# Loop Extrusion Mediates Physiological Locus Contraction for V(D)J Recombination

**DOI:** 10.1101/2020.06.30.181222

**Authors:** Hai-Qiang Dai, Hongli Hu, Jiangman Lou, Adam Yongxin Ye, Aimee M. Chapdelaine-Williams, Nia Kyritsis, Yiwen Zhang, Nicole Manfredonia, Rachael Judson, Huan Chen, Kerstin Johnson, Sherry Lin, Zhaoqing Ba, Andrea Conte, Rafael Casellas, Cheng-Sheng Lee, Frederick W. Alt

## Abstract

Immunoglobulin heavy chain locus (*Igh*) V_H_, D, and J_H_ gene segments are developmentally assembled into V(D)J exons. RAG endonuclease initiates V(D)J recombination by binding a J_H_-recombination signal sequence (RSS) within a chromatin-based recombination center (RC) and then, in an orientation-dependent process, scans upstream D-containing chromatin presented by cohesin-mediated loop extrusion for convergent D-RSSs to initiate DJ_H_-RC formation^1,2^. In primary pro-B cells, 100s of upstream V_H_-associated RSSs, embedded in convergent orientation to the DJ_H_-RC-RSS, gain proximity to the DJ_H_-RC for V_H_-to-DJ_H_ joining via a mechanistically-undefined V_H_-locus contraction process^3-7^. Here, we report that a 2.4 mega-base V_H_ locus inversion in primary pro-B cells nearly abrogates rearrangements of normally convergent V_H_-RSSs and cryptic RSSs, even though locus contraction *per se* is maintained. Moreover, this inversion activated rearrangement of both cryptic V_H_-locus RSSs normally in the opposite orientation and, unexpectedly, of normally-oriented cryptic RSSs within multiple, sequential upstream convergent-CBE domains. Primary pro-B cells had significantly reduced transcription of *Wapl*^8^, a cohesin-unloading factor, versus levels in *v-Abl* pro-B lines that lack marked locus contraction or distal V_H_ rearrangements^2,9-11^. Correspondingly, Wapl depletion in *v-Abl* lines activated V_H_-locus contraction and orientation-specific RAG-scanning across the V_H_-locus. Our findings indicate that locus contraction and physiological V_H_-to-DJ_H_ joining both are regulated via circumvention of CBE scanning impediments.

---

The ability of dominant CBE contact loops over mega-base (Mb) distances genome-wide to be anchored predominantly by convergently-oriented CBEs versus CBEs in other orientations was proposed to be mediated by cohesin-mediated loop extrusion^12,13^, as opposed to potential diffusion-based mechanisms that cannot readily explain orientation-bias. Parallels between findings on RAG chromatin exploration during V(D)J recombination and loop extrusion models provided a foundation for the discovery of fundamental roles of loop extrusion in D to J_H_ recombination^1,9,10,12-14^. Indeed, while Ds have downstream and upstream RSSs in opposite orientations, only downstream D-RSSs convergently-oriented with initiating J_H_-RSSs are used for D to J_H_ joining^1^. However, while all V_H_-RSSs are in convergent orientation with the upstream DJ_H_-RC-RSS^9^, potential access of the upstream V_H_s, at the D-to-J_H_ rearrangement stage, via linear scanning is largely blocked by CBEs in intergenic control region 1 (IGCR1) between the V_H_ and D loci^9-11^. In this regard, V_H_ utilization across the 2.4 Mb V_H_ locus has been associated with physical *Igh* locus contraction in pro-B cells, which is proposed to bring V_H_s into close orbit around the DJ_H_-RC to allow V_H_s to gain RC access via diffusion^3-7^. In its simplest form, such a diffusion-based model also predicts that V_H_-RSS orientation would not play a dominant role in V_H_ utilization across the V_H_ locus.

As an initial test of a diffusion-based versus linear scanning model for long-range V_H_ utilization, we inverted the entire 2.4Mb V_H_ locus upstream of the D-proximal V_H_81X in the genome of C57BL/6 mice,leaving V_H_81X in normal position and orientation as a control (Fig. 1a, Extended Data Fig. 1). In C57BL/6 mice, the entire V_H_ locus has been sequenced, revealing 109 functional V_H_s with locations accurately mapped within four V_H_ domains referred to from proximal to distal as: 7183/Q52, Middle, J558, and J558/3609, respectively^15,16^ (Fig. 1a, top, Supplementary Table 1). This large scale inversion placed the distal J558/3609 V_H_s in the proximal V_H_ location and the proximal 7183/Q52 V_H_ locus, besides V_H_81X, in the distal location, with all 108 functional V_H_s and their RSSs across the locus now in inverted orientation relative to the DJ_H_-RC RSS (Fig. 1a middle, bottom). Based on our sensitive HTGTS-V(D)J-seq assay^17,18^, this inversion nearly abrogated rearrangements of the normally convergent V_H_-RSSs across the inverted region in primary bone marrow (BM) pro-B cells, with utilization of the normally oriented and positioned proximal V_H_81X actually doubling, perhaps due to lack of competition from immediately upstream V_H_s^10^ (Fig. 1b, top versus bottom panel; Supplementary Table 1).

**Figure 1.**
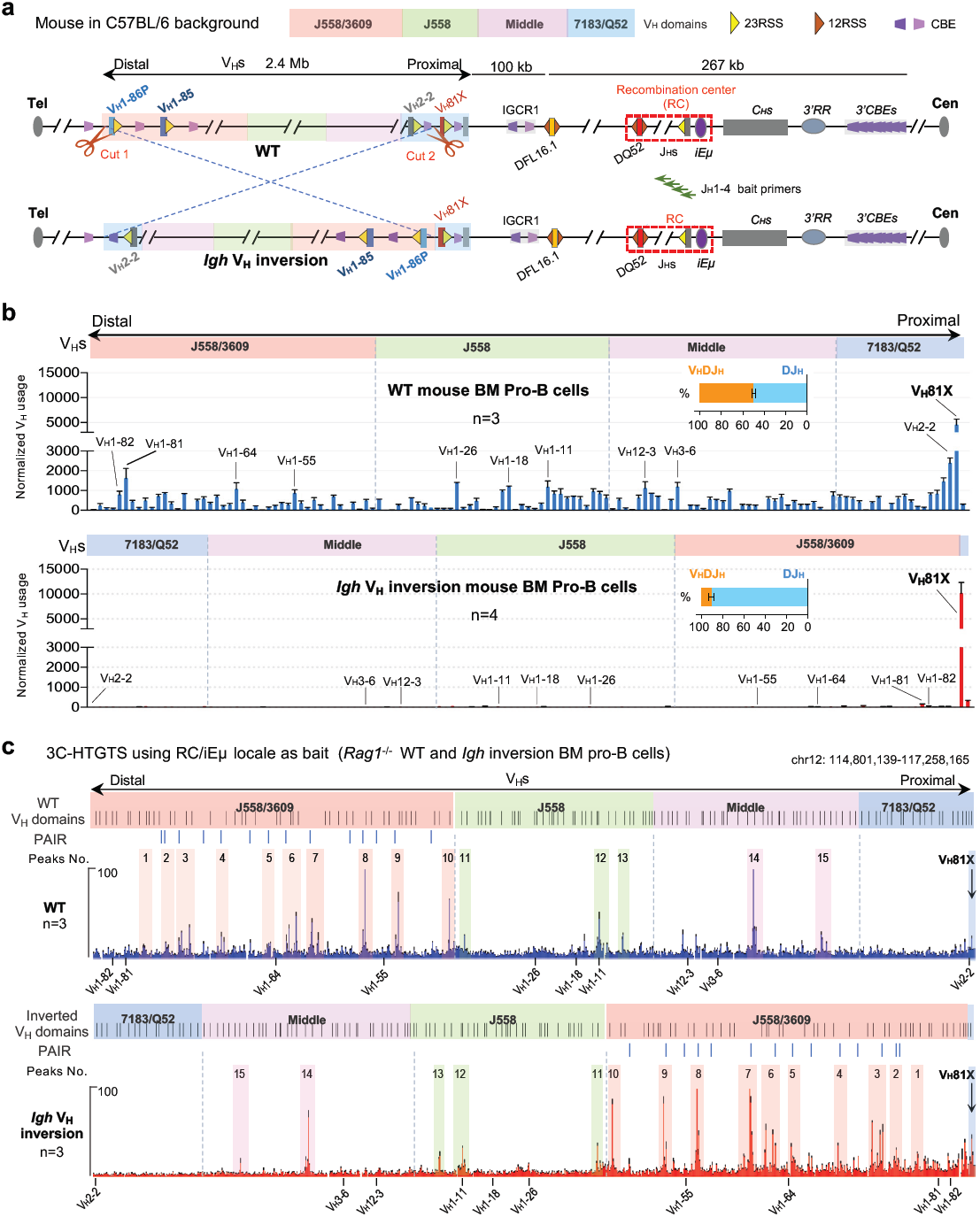
*Igh* V_H_ inversion mutation marked decreased utilization of all V_H_s other than the still convergently oriented V_H_81X in primary BM pro-B cells. **a**, Schematic of the strategy to generate the mouse model with entire *Igh* V_H_ locus inversion upstream V_H_81X. Diagram of the murine *Igh* locus showing the first two proximal V_H_s (V_H_81X and V_H_2-2), the last two distal V_H_s (V_H_1-86P and V_H_1-85), Ds, J_H_s, C_H_s, and regulatory elements as indicated (not to scale), with RC that comprises J_H_-proximal DQ52 segment, four J_H_ segments, and the intronic enhancer (iEμ) highlighted. All V_H_ segments divided into four V_H_s domains (7183/Q52, Middle, J558 and J558/3609) from most proximal to distal. Yellow and dark orange triangles represent position and orientation of *bona fide* 23RSS and 12RSS, respectively. Purple and pink trapezoids represent position and orientation of CBEs. Green arrows denote the J_H_1-4 coding end bait primers used for generating HTGTS-V(D)J-seq libraries. Cut1 and Cut2 showing the location of 2 sgRNAs. Tel, telomere. Cen, centromere. **b**, Average utilization frequencies ± s.d. of all V_H_ segments in WT (top, blue, n=3) and *Igh* V_H_ inversion (bottom, red, *n*=4) mouse BM pro-B cells. Average percentage ± s.d. of V_H_DJ_H_ and DJ_H_ rearrangements are indicated. For comparison, several highly utilized V_H_s in each V_H_ domain are indicated. See Supplementary Table 1 and Methods for more details. **c**, Average 3C-HTGTS signal counts ± s.e.m. across the four V_H_ domains in cultured WT (top, n=3) and *Igh* V_H_ inversion (bottom, n=3) *Rag1*^*-/-*^ mouse BM pro-B cells. The WT and inverted V_H_ locus/domains with PAIR elements are diagrammed at the top of each panel. For comparison, the location of the highly utilized V_H_s in (**b**) are labeled below each panel (**c**). 15 representative major interaction peaks/clusters were indicated with colour shades and numbers. See Extended Data Fig. 2 and Supplementary Data 1 for more details. In all panels, “n” indicates biological repeats. *p* values were calculated using unpaired two-tailed t-test.

To examine effects of the inversion on interactions of V_H_ locus sequences with the RC, we performed high-resolution 3C-HTGTS^10^ with a RC bait in RAG1-deficient cultured BM pro-B cells with or without the inversion. RAG-deficient cells were used to prevent confounding V(D)J recombination events^10^. These studies revealed a series of robust peaks of RC interaction across the V_H_ locus (Fig. 1c, labeled 1-15), a number of which, including those involving Pax5-activated intergenic repeat (PAIR) elements^19^ (Fig. 1c, blue bars), are considered hallmarks of V_H_ locus contraction^5^. Notably, all of the major RC-V_H_ locus interaction peaks were maintained within the inverted V_H_ locus, despite the huge positional exchanges between more proximal and more distal portions of the locus (Fig. 1c; labeled 1-15; Extended Data Fig. 2a-d). Indeed, the overall RC interactions within the different regions of the inverted locus appeared almost as a mirror-image of the patterns of the normally-oriented locus (Fig. 1c, upper and lower). Thus, the 2.4Mb inversion did not markedly disrupt V_H_-locus contraction *per se.* In addition, cohesin binding patterns determined by ChIP-seq and germline V_H_ transcription patterns determined by GRO-seq across the different V_H_ domains were recapitulated in detail in the inverted V_H_-locus, again largely as mirror images of those of the normal V_H_-locus (Extended Data Fig. 3a-d; Supplementary Data 1). Also notable is that many of the major interaction peaks corresponded to high level peaks of sense/antisense transcription, although within the inverted V_H_ locus orientation of sense and antisense transcription relative to the RC is reversed (Extended Data Fig. 3d, bottom). However, despite maintenance of RC interactions, transcription patterns, and binding of key looping factors (Fig. 1c; Extended Data Fig. 2e and 3a-d; Supplementary Data 1), the inverted V_H_-RSSs were barely used by the V(D)J recombination process. This finding is consistent with the notion that DJ_H_-RC-bound RAG linear scanning of V_H_ locus chromatin plays an important role in V_H_ utilization in V_H_-locus contracted BM pro-B cells.

It remained conceivable that the large-scale inversion could have somehow prevented V_H_s across the inverted locus from accessing the DJ_H_-RC via diffusion following locus contraction. Introduction of ectopic RCs into non-antigen receptor loci across the genome in *v-Abl* pro-B lines allows RAG, in the absence of other major scanning impediments, to linearly explore convergent CBE-anchored chromatin domains, up to at least several Mb distances, until scanning is terminated at convergent CBE loop anchors^9^. In these experiments, orientation of initiating *bona fide* RSSs programs RAG activity linearly in one direction or the other, up to Mb distances, with linear scanning evidenced by low-level of utilization 100s of cryptic RSSs, as short as the conserved first 3 bp of the RSSs (CAC), but only when they are in convergent orientation with the initiating *bona fide* RSS^9^. In such experiments, cryptic RSSs in the same orientation as the RC-RSS (e.g. GTGs) are not used, demonstrating that cryptic RSS access ectopic RCs by linear scanning versus diffusion^9^. Thus, to further explore the predictions of a linear RAG scanning model for V_H_ utilization in BM pro-B cells, we analyzed RAG utilization of cryptic RSSs across the V_H_ locus in BM pro-B cells (Fig. 2a, top). These analyses clearly demonstrated that cryptic RSSs across the length of 2.4 Mb V_H_ locus in convergent orientation with the DJ_H_-RC-RSS were utilized, but those in the same orientation as the DJ_H_-RC-RSS were not, strongly indicating that RC-bound RAG linearly scans across the V_H_ locus in BM pro-B cells (Fig. 2b, top and 2c; Extended Data Fig.4a, top, and 4b). This analysis also revealed that cryptic RSS utilization did not extend far beyond the *Igh* locus, likely due to scanning activity largely being terminated within the V_H_ locus in most BM pro-B cells by robust rearrangement of V_H_s with convergent *bona fide* RSSs (Fig.1b, top; Fig. 2b, top and 2c; Extended Data Fig. 4a, top, and 4b).

**Figure 2.**
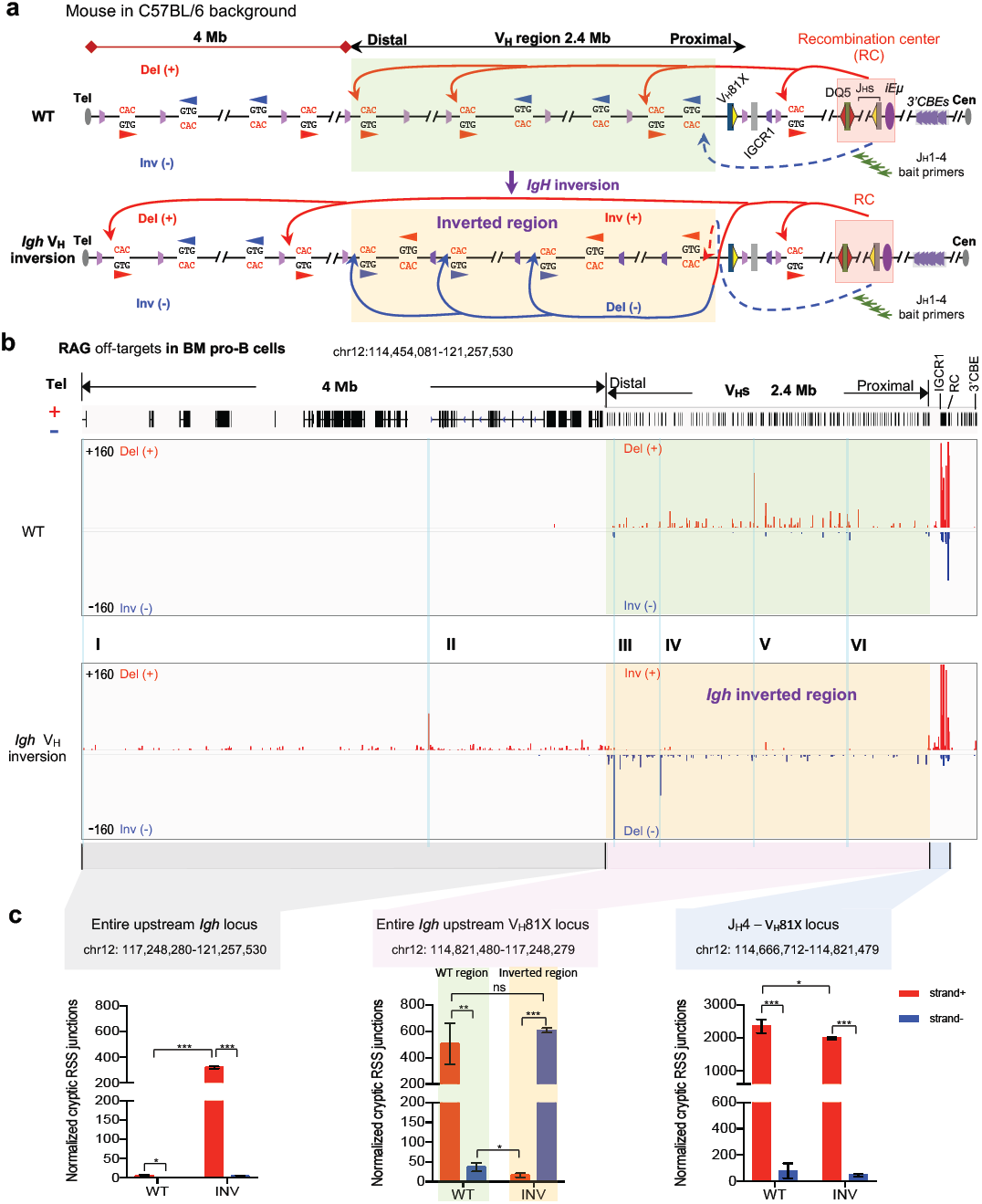
RAG targets cryptic RSSs that *Igh* locus inversion place in convergent orientation to the upstream D-RSSs in primary BM pro-B cells. **a**, Illustration of possible joining outcomes between *bona fide* RSSs from J_H_1-4 coding end baits to cryptic RSSs mostly represented by CAC motifs in the upstream Ds, V_H_s and upstream region till telomere in WT (top) and *Igh* V_H_ inversion (bottom) mouse primary pro-B cells. Red and blue arches with arrows show possible deletional and inversional junctions, respectively, except for the *Igh* V_H_ inverted region (red for inversional, blue for deletional). Green arrows indicate the position and orientation of HTGTS primers. Details as shown in Fig. 1a. **b**, RAG off-target junction profiles at *Igh* locus and upstream 4 Mb region in WT (middle) and *Igh* V_H_ inversion (bottom) BM pro-B cells isolated from individual mice. Top panel is WT 2.4 Mb *Igh* locus and upstream 4 Mb region track. Junctions are displayed with linear scale as stacked tracks. I-VI, indicated by sky blue lines, are examples of RAG off-target peaks plotted at single-base-pair resolution in Extended Data Fig. 4a. The background of *Igh* V_H_ inverted region is highlighted in transparent green (**a, b**, WT) and transparent gold (**a, b**, *Igh* V_H_ inversion), respectively. Del (+), deletional junction. Inv (-), inversional junction. Each library was normalized. See methods and related Extended Data Fig. 4 for more details and additional independent repeats. **c**, Average frequencies ± s.d. of plus strand (red, +) and minus strand (blue, -) joining events within indicated regions in wide-type (WT, n=3) and *Igh* V_H_ inversion (INV, *n*=3) mouse BM pro-B cells. Three indicated regions (J_H_4-V_H_81X locus, Entire *Igh* upstream V_H_81X locus, and Entire upstream *Igh* locus) were highlighted (**b**), respectively. Each of the biological library replicates was normalized to 2979 off-target junctions for statistical analysis. *p* values were calculated using unpaired two-tailed t-test. See Methods for more details.

The large V_H_ inversion locus provided the opportunity for an even more stringent test of the linear scanning model. Thus, the scanning model predicts within the inverted V_H_ locus, cryptic RSSs that normally were not utilized because they were in the same orientation as the DJ_H_-RC-RSS would now be more robustly utilized due to being placed in convergent orientation by the inversion (Fig. 2a, bottom). Indeed, the 2.4Mb V_H_-locus inversion essentially eliminated utilization of the previously convergently-oriented cryptic RSSs now placed in the same orientation as the DJ_H_-RC-RSSs and activated utilization of cryptic RSSs previously in the same orientation that were now placed in convergent orientation (Fig. 2b, bottom and 2c; Extended Data Fig.4a, bottom, 4c). Strikingly, the inversion also resulted in RAG scanning well beyond the end of the *Igh* locus in BM pro-B cells over distances of 4 Mb upstream, nearly to the telomere (Fig. 2b, bottom and 2c; Extended Data Fig. 4a, bottom, 4c). Moreover, consistent with linear RAG scanning through the long non-inverted region upstream of the V_H_ locus, cryptic RSS utilization indeed involved those normally in convergent orientation with the DJ_H_-RC-RSS (Fig.2b, bottom and 2c; Extended Data Fig. 4a, bottom, 4c). This continuation of RAG scanning upstream of the V_H_ locus in the context of the V_H_ locus inversion suggests that substantial inactivation of dominant V_H_ rearrangements, due to their inversion, allows DJ_H_-RC-initiated RAG scanning to more robustly proceed through the V_H_ locus into the upstream region. RAG scanning through the CBE-rich *Igh* locus in BM pro-B cells with normal or inverted *Igh* loci, as well as RAG scanning through multiple upstream convergent CBE-based domains, which are well-known to impede scanning^9,20-22^, suggests that BM pro-B cells may have an active mechanism to neutralize ability of V_H_-associated CBEs to impede RAG scanning that influences convergent CBE domains more broadly than those just within the V_H_ locus.

G1-arrested *v-Abl* pro-B cells can be viably arrested in the G1 cell-cycle stage by treatment with *Abl* tyrosine kinase inhibitor (STI-571)^23^, during which time they induce RAG expression and undergo robust D to J_H_ joining, but very little V_H_-to-DJ_H_ joining in association with little V_H_-locus contraction^1,10^. Thus, as one approach to elucidate a potential CBE-neutralizing mechanism in normal BM pro-B cells, we employed GRO-seq to measure the transcriptional activity of various genes encoding proteins implicated in cohesin complex function. We did this in locus-contracted RAG1-deficient C57BL/6 mouse BM pro-B cells and G1-arrested *v-Abl*-transformed *Eμ-Bcl2*-expressing pro-B cells (“*v-Abl* pro-B cells”), which were also derived from RAG1-deficient C57BL/6 mouse BM pro-B cells (Extended Data Fig. 5a). Notably, of all factors analyzed, we found only the Wings apart-like (*Wapl*) factor^8^ to have very low transcription in BM pro-B cells versus *v-Abl* pro-B cells (Fig. 3a, b). Wapl is known to unload cohesin from chromosomes by transiently opening the cohesin ring^24-26^ and, thereby, contributes to regulation of chromatin structure and protects against chromosome segregation errors and aneuploidy^27,28^. Cohesin also is positioned in genomes by transcription, CTCF and Wapl^29,30^. Most relevantly, Wapl-depletion extends CTCF-anchored loop sizes in mammalian cells in association with increasing cohesin density^31^, which may increase the probability of extending loop extrusion past dynamic CTCF-bound CBE loop anchors^32,33^. Based on the finding of relatively low *Wapl* transcription in BM pro-B cells, we tested effects of substantial Wapl-depletion on *Igh* loop domain formation and RAG scanning in G1-arrested C57BL/6 mouse *v-Abl* pro-B cells.

**Figure 3.**
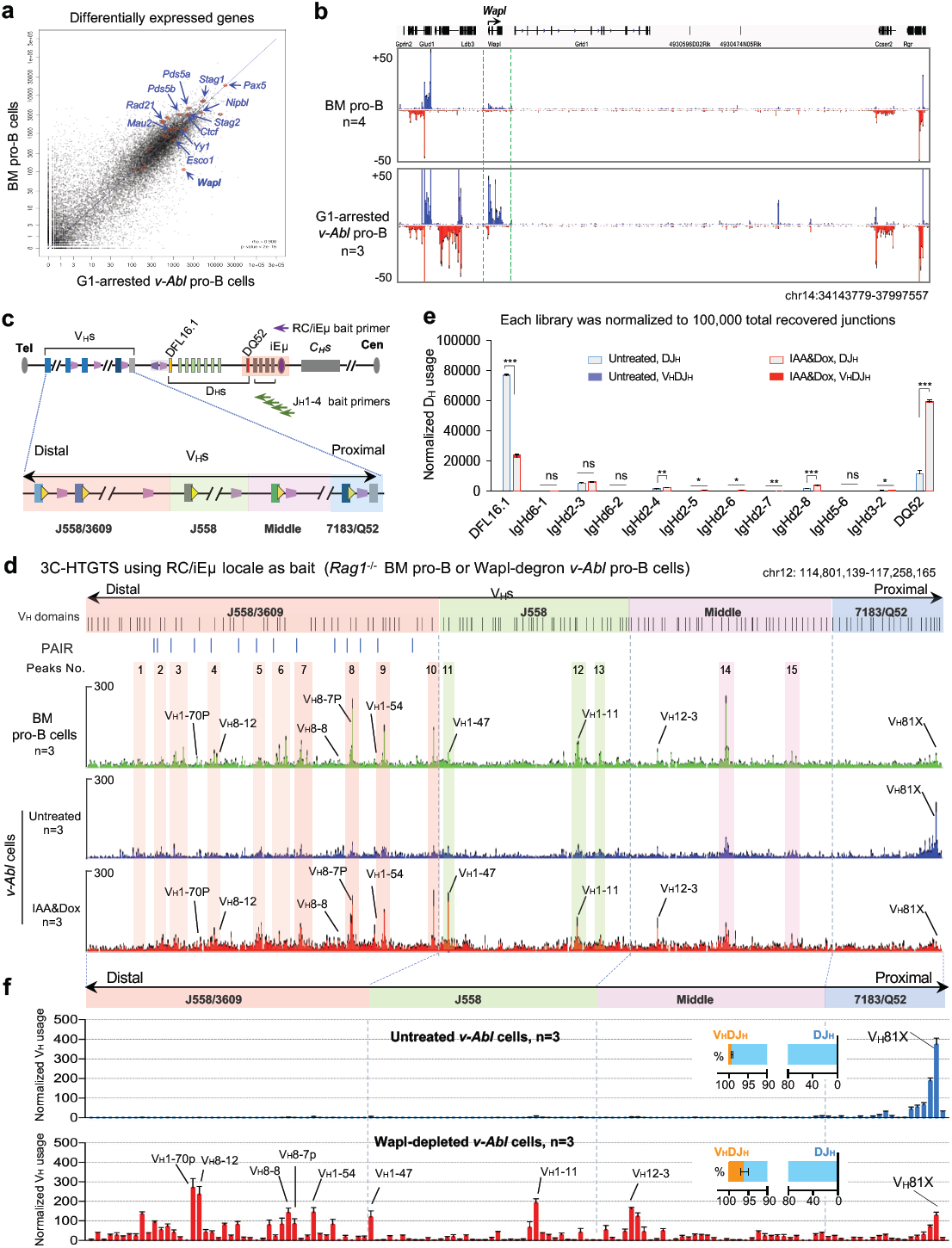
Wapl depletion activates *Igh* RC interactions with the V_H_ locus and results in increased distal V_H_s utilization in G1-arrested Wapl-degron *v-Abl* pro-B cells. **a**, Scatter plots of averaging transcriptome-wide GRO-seq counts in G1-arrested *v-Abl* pro-B cells (*x axis*, n=3) and BM pro-B cells (*y axis*, n=4). Representative known requisite genes implicated in the cohesin complex function for V(D)J recombination and chromatin interaction are highlighted by red circles and blue arrows. Spearman’s correlation coefficient (rho) and *p* value determined by two-sided Spearman’s correlation test are presented. **b**, Comparation of *Wapl* transcription level in BM pro-B cells and G1-arrested *v-Abl* pro-B cells from 4 and 3 independent repeats, respectively. The transcription level of genes lying upstream and downstream of *Wapl* were shown in the same panel. **c**, Diagram of the entire murine *Igh* locus with details as shown in Fig. 1a. Purple and green arrows denote the RC/iEμ bait and J_H_1-4 coding end bait primers used for 3C-HTGTS and HTGTS-V(D)J-seq, respectively. **d**, Average 3C-HTGTS signal counts ± s.e.m. across the four V_H_ domains in cultured *Rag1*^*-/-*^ BM pro-B cells (top, n=3), G1-arrested *Rag1*^*-/-*^ *v-Abl* pro-B cells without (middle, n=3) or with Wapl depletion (bottom, n=3). For comparison, 15 representative major interaction peaks/clusters as shown in Fig.1c were indicated with colour shades and numbers. PAIR elements are indicated as blue bars. **e, f**, Average utilization frequencies ± s.d. of all D segments (**e**) from DJ_H_ and V_H_DJ_H_ joins, and all V_H_ segments (**f**) in RAG1-complemented, G1-arrested Wapl-degron *v-Abl* pro-B cells without (Untreated, n=3) or with (IAA&Dox, red, n=3) Wapl depletion. Average percentage ± s.d. of V_H_DJ_H_ and DJ_H_ rearrangements are indicated (**f**). For comparison, several highly utilized V_H_s (**f)** in Wapl-depleted *v-Abl* pro-B cells and the corelated interaction peaks (**d**) are highlighted, respectively. *p* values were calculated using unpaired two-tailed t-test. See Supplementary Table 2 and Methods for more details. Notably, while Wapl-depleted *v-Abl* cells have RC interactions across the V_H_ locus that are very similar to those of BM pro-B cells (**d**, upper and bottom); their V_H_ to DJ_H_ rearrangement levels are much lower than those of BM pro-B cells (compare Fig. 1b and 3f), despite having activated RAG scanning across the entire V_H_ locus (**d**). These observations may at least in part reflect RC-interactions being assayed in RAG-deficient cells while V(D)J recombination is assayed in RAG sufficient cells. In this regard, prior studies suggested extrusion of chromatin impediments past the RC may be more efficient when it is not RAG-bound^10^.

We generated a RAG1-deficient C57BL/6 *v-Abl* pro-B cell line that expressed the TIR1 F-box protein from *Oryza sativa* (OsTIR1) under control of the conditional Tet promoter (Tet-on-OsTIR1)^34^. OsTIR1 binds a mini auxin-inducible degron (mAID) in the presence of auxin to trigger proteasome-dependent degradation^34^. We refer to these modified cells as the “primary” *v-Abl* line (Extended Data Fig. 5b, c). We subsequently targeted a stop codon into both *Wapl* alleles in the primary *v-Abl* line to introduce in-frame sequences encoding mAID with a fluorescent monomeric Clover (mClover) cassette^34^ (Extended Data Fig. 5d, e), and refer to the resulting lines as “Wapl-degron *v-Abl* pro-B cells”. For experiments, we added doxycycline (Dox) and auxin (Indole-3-acetic acid, IAA) to induce OsTIR1 expression and trigger proteasome-dependent degradation of the Wapl-mAID-mClover protein, respectively. To specifically deplete mAID-tagged Wapl protein in the G1 stage, we treated the Wapl-degron *v-Abl* pro-B cells with IAA and Dox from day1 to day4 during the STI-571-induced four-day G1 arrest experimental period (day0-4) (Extended Data Fig. 5f). Western blotting revealed that mAID-tagged Wapl was substantially depleted in G1-arrested IAA/Dox-treated, but not untreated, Wapl-degron *v-Abl* pro-B cells (Extended Data Fig. 5g, h). While Wapl depletion is lethal to dividing cells^27,28^, Wapl-depleted *v-Abl* pro-B cells underwent normal STI-571-induced G1 arrest with only modest effects on viability over the remaining experimental period (Extended Data Fig. 5i, j). Substantial depletion of chromatin-bound mAID-tagged Wapl across the *Igh* locus and genome-wide in G1-arrested Wapl-depleted *v-Abl* cells was confirmed by ChIP-seq (Extended Data Fig. 6a, b). Robust sense and anti-sense transcription through the RC, genome-wide transcription patterns, and binding of several key looping factors across V_H_ domains were maintained in G1-arrested, IAA/Dox-treated Wapl-degron *v-Abl* cells (Extended Data Fig. 6c-e).

3C-HTGTS-based interaction experiments of untreated versus IAA/Dox-treated G1-arrested RAG1-deficient Wapl-degron *v-Abl* pro-B cells revealed low-level peaks of RC interactions in all but the proximal 7183/Q52 domain V_H_ locus in untreated cells that were substantially increased upon Wapl depletion (Fig. 3c, d middle and bottom). Indeed, the major interactions peaks across the J558/3609, J558, and Middle V_H_ domains that were substantially increased in IAA/Dox-treated Wapl-degron *v-Abl* pro-B cells closely corresponded to the dominant peaks observed in the context of locus-contracted RAG1-deficient, cultured BM pro-B cells (Fig. 3d, upper, labeled 1-15), indicating that Wapl-depletion in *v-Abl* lines enhanced V_H_-locus contraction (Fig. 3d). This finding supports the notion that locus contraction may, at least in part, involve modulation of ability of cohesin-mediated loop extrusion to circumvent CTCF-bound CBE impediments. Baits specific for either a prominent V_H_J558/3609 interaction peak or a prominent V_H_J558 interaction peak locale showed increased interactions with the RC upon Wapl-depletion in the absence of increased interactions with other *Igh* locales (Extended Data Fig. 7a, b). If extrapolated to be representative RC interactions of all separate V_H_ locus interaction sites across the V_H_ domains in Wapl depleted *v-Abl* lines, these latter findings predict that a substantial fraction of individual cells in the population would have one or another sequence across their V_H_ locus interacting with the RC.

To test effects of Wapl-depletion on *Igh* V(D)J recombination, we introduced RAG1 into RAG1-defcient C57BL/6 Wapl-degron cells (Extended Data Fig. 8a). Similar to primary *v-Abl* cells (Extended Data Fig. 8b, c), untreated Wapl degron *v-Abl* pro-B cells activated robust D to J_H_ rearrangement with J_H_-distal DFL16.1 having highest rearrangement frequency and J_H_-proximal DQ52 intermediate frequency (Fig. 3e, Supplementary Table 2). As anticipated, untreated cells had very little V_H_ to DJ_H_ rearrangement, with the vast majority of that which did occur involving the V_H_-CBE dependent rearrangement of the very most proximal V_H_s^1,10^ (Fig. 3f, upper; Extended Data Fig. 6e, upper). Wapl depletion markedly reduced distal DFL16.1 rearrangement levels, while promoting greatly increased proximal DQ52 rearrangement (Fig. 3e; Supplementary Table 2), a pattern also observed in the context of inactivation of IGCR1 CBEs^10,35^ (Extended Data Fig. 8b, e, f; Supplementary Table 3). Strikingly, Wapl depletion promoted an approximately four-fold overall increased V_H_ to DJ_H_ recombination, associated with significantly increased utilization of V_H_s across the four V_H_ domains of the 2.4 Mb *Igh* locus (Fig. 3f, bottom; Supplementary Table 2). However, an exception was that utilization of the very most proximal V_H_s was significantly decreased (Fig. 3f, bottom; Supplementary Table 2), which is consistent with diminished impediment activity of the most proximal V_H_-CBEs^10^ (Extended Data Fig. 6e, upper; Extended Data Fig. 8d, bottom). Together, the interaction and V(D)J recombination data are consistent with Wapl extending loop extrusion past CTCF-bound IGCR1 CBEs and proximal V_H_-CBEs. To further assess this apparently enhanced long-range RAG scanning, we analyzed RAG utilization of cryptic RSSs across the V_H_ locus in Wapl-depleted *v-Abl* pro-B cells. Consistent with the findings in locus-contracted BM pro-B cells (Fig. 2b, upper), in Wapl depleted *v-Abl* pro-B cells, RAG utilized cryptic RSSs across the length of 2.4 Mb V_H_ locus that were in convergence with, but not in the same orientation as, the initiating DJ_H_-RC-RSS for joining (Extended Data Fig. 9). Together, these studies demonstrate that Wapl depletion facilitates orientation-specific scanning of RC-bound RAG across the entire V_H_ locus.

We demonstrate that RAG accesses most V_H_s across the 2.4 Mb V_H_ locus in BM Pro-B cells by a linear, orientation-dependent scanning mechanism. Our finding of RC-based RAG scanning through multiple convergent CBE domains upstream of an inverted V_H_ locus suggests that pro-B cells may activate V_H_-locus scanning via a mechanism that allows loop extrusion to bypass, at least partially, CTCF-bound CBEs more generally. Very low *Wapl* transcription in BM pro-B cells led us to deplete Wapl in *v-Abl* cells, which markedly enhanced both V_H_ locus contraction and long-range V_H_ utilization, indicating that both processes involve cohesin-mediated loop extrusion in the context of diminished CBE impediments. In this regard, our detailed analyses of individual V_H_s are in accord with recent CTCF-depletion studies in *v-Abl* lines^2^ that show residual CBE activity and/or transcription may enhance utilization of many V_H_s along the scanning path (Supplementary Data 2). While RAG has been shown to scan directly over Mb distances across genomic sequences that lack major loop extrusion impediments^9^, other mechanisms for RAG access of distal V_H_s by linear, orientation-dependent scanning of the impediment rich V_H_ locus are conceivable. For example, loop extrusion-mediated locus contraction may occur frequently in BM pro-B cells when RAG is not bound to the RC. In such cells, partial neutralization of CBE impediments could allow loop extrusion past the RC to proceed for varying distances across the locus (Fig. 3, legend). Then, upon RAG-binding to the RC, linear scanning could be initiated at the point extrusion has reached along the locus in a given pro-B cell. In this way, loop extrusion mediated-locus contraction could augment ability of RAG scanning to access more distal V_H_s. Finally, our Wapl-depletion findings show that, beyond modulating CTCF activity^2^, V_H_ locus CBE neutralization can occur by modulating cohesin ability to circumvent CTCF-bound CBEs.

## ACKNOWLEDGEMENTS

We thank lab members for stimulating discussions, Hwei-Ling Cheng for advice and help with ES cell culture, Ming Tian for EF1 ES cell line, Yu Zhang and Xuefei Zhang for RAG-expressing retrovirus plasmids. This work was supported by NIH R01 AI020047 (to F.W.A.). H.-Q.D. is a fellow of the Cancer Research Institute (CRI) of New York. H.C. is a NRSA Fellow (T32 ai07386) and was supported by a Leukemia and Lymphoma Society fellowship. Z.B. was supported by CRI fellowship. R.C. and A.C. are partially funded by the NIH Regulome Project. C.-S.L. was previously supported by CRI fellowship and is now funded by the Ministry of Science and Technology in Taiwan [MOST109-2636-B-007-004]. F.W.A. is an investigator of the Howard Hughes Medical Institute.

## AUTHOR CONTRIBUTATIONS

H.-Q.D., C.-S.L. and F.W.A. conceived the original idea and designed the study. H.-Q.D. generated and analysed the *Igh* V_H_ locus inversion mouse model, and found the significantly low-level of *Wapl* transcription in BM pro-B cells. C.-S.L. initiated and established the Wapl-degron *v-Abl* system and obtained preliminary V usage and interaction results, and together with H.-Q.D. generated the doxycycline inducible Wapl-degron lines. H.-Q.D. optimized the RAG complementation method in the Wapl-degron system and performed the analyses, and together with H.L. and C.-S.L. performed 3C-HTGTS. H.L. performed ChIP-seq, Western blot and Southern blot. J.L. performed GRO-seq. A.Y.Y. and N.K. designed some of the bioinformatics pipelines for data analyses. A.M.C.-W with the assistance of N.M., R.J. and K.J. performed RDBC injections. H.C. optimized HTGTS-V(D)J-seq. Y.Z. assisted with the analysis of data. Z.B. and S.L established the RAG1-deficient, *Eμ-Bcl2*-expressing C57BL/6 *v-Abl* pro-B cell line. H.-Q.D., H.L. and F.W.A., with input from C.-S.L., A.Y.Y., analyzed and interpreted data, designed figures, and wrote the manuscript. C.-S.L., R.C., J.L., Y.Z., Z.B. and A.C. provided insights and helped polish the paper. F.W.A. supervised the study.

## AUTHOR INFORMATION

The authors declare no competing financial interests. Correspondence and requests for materials should be addressed to F.W.A. F.W.A. is a co-founder of Otoro Biopharmaceuticals.

## EXTENDED DATA FIGURE LEGENDS

**Extended Data Figure 1.**
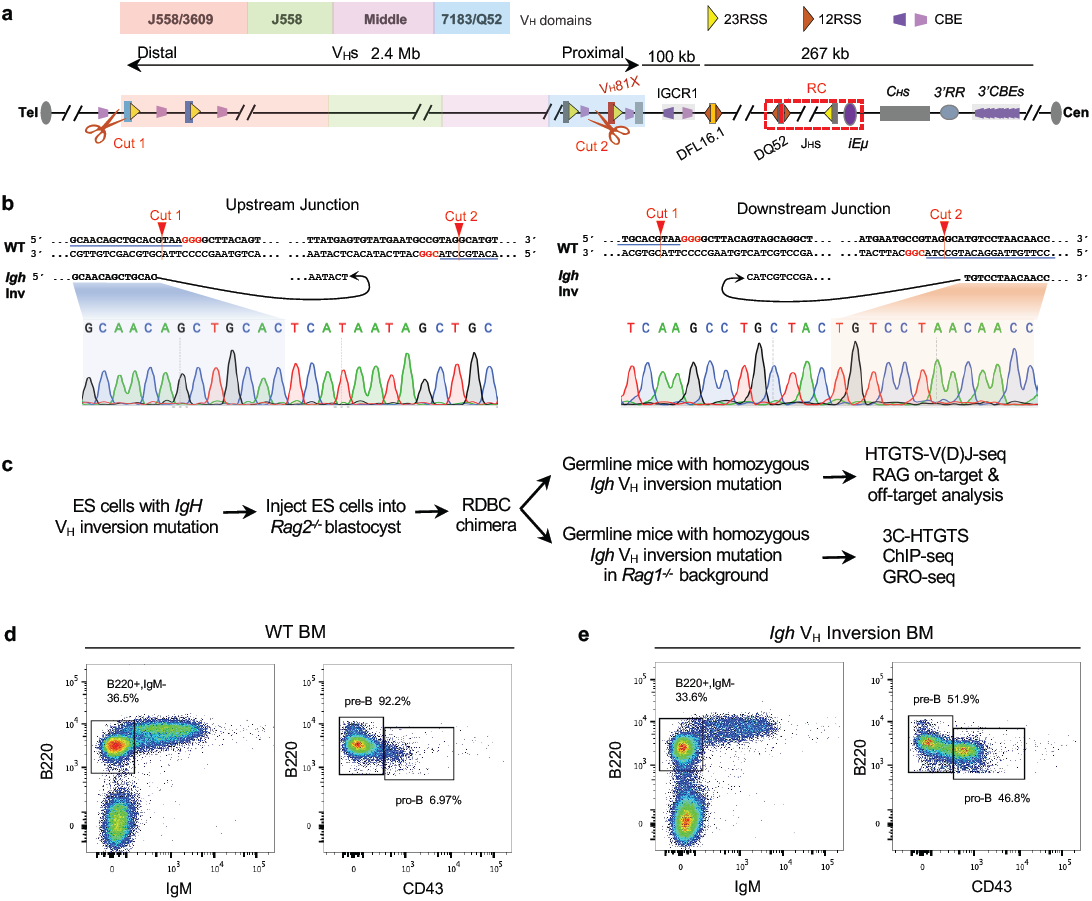
Generation and characterization of *Igh* V_H_ locus inversion mouse model. **a**, Schematic diagram showing CRISPR-Cas9-mediated entire *Igh* V_H_ locus inversion upstream V_H_81X in embryonic stem cells (ES cells) on the *Igh* allele in C57BL/6 genetic background. Cut1 and Cut2 showing the location of 2 sgRNAs. Details as shown in Fig. 1a. **b**, Confirmation of the upstream and downstream inversion junctions by Sanger sequencing. The sgRNA-targeting sequence is underlined, and the PAM sequence is labelled in red. sgRNAs and oligos used are listed in Supplementary Table 4. **c**, Schematic showing the generation of *Igh* V_H_ locus inversion mouse model and further assays for phenotype and mechanism analysis. **d, e**, Representative flow cytometry analysis of IgM^-^ bone marrow (BM) B cell populations in 4∼6-week-old WT (**d**) and *Igh* V_H_ locus inversion (**e**) mice. B220^+^IgM^-^ B cells were gated and shown in the left plot (**d, e**). B220^+^CD43^+^ pro-B and B220^+^CD43^-^ pre-B cell populations are indicated in the right plot (**d, e**).

**Extended Data Figure 2.**
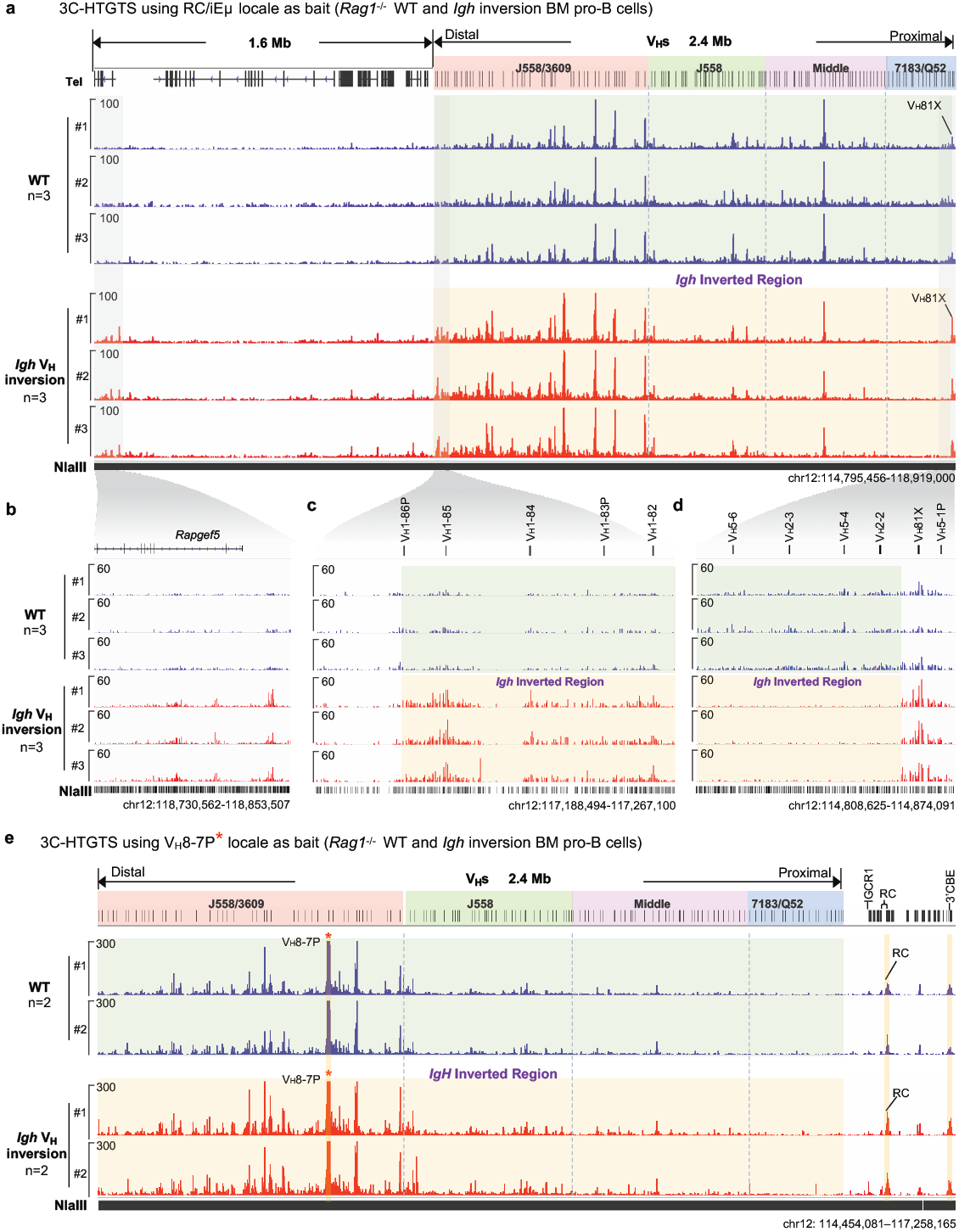
3C-HTGTS chromatin interaction profiles at *Igh* locus and upstream region in WT and *Igh* V_H_ locus inversion cultured *Rag1*^*-/-*^ BM pro-B cells. For comparison, all the data were shown in normal *Igh* orientation. **a**, 3C-HTGTS chromatin interaction profiles of RC/iEμ bait across the 4 V_H_ domains and 1.6 Mb upstream region in WT (blue, n=3) and *Igh* V_H_ locus inversion (red, *n*=3) cultured *Rag1*^*-/-*^ mouse BM pro-B cells. **b-d**, 3C-HTGTS profiles zoomed in the upstream *Igh* region (**b**), distal *Igh* V_H_ region (**c**), and proximal *Igh* V_H_ region (**d**). **e**, 3C-HTGTS chromatin interaction profiles of distal V_H_8-7P bait across the entire *Igh* locus in WT (blue, n=2) and *Igh* V_H_ locus inversion (red, *n*=2) cultured *Rag1*^*-/-*^ mouse BM pro-B cells. All V_H_ segments divided into four V_H_s domains (7183/Q52, Middle, J558 and J558/3609) from most proximal to distal. The background of *Igh* V_H_ inverted region is highlighted in transparent green (WT) and transparent gold (*Igh* V_H_ inversion), respectively. In all panels, “n” indicates biological repeats. See Methods for more details.

**Extended Data Figure 3.**
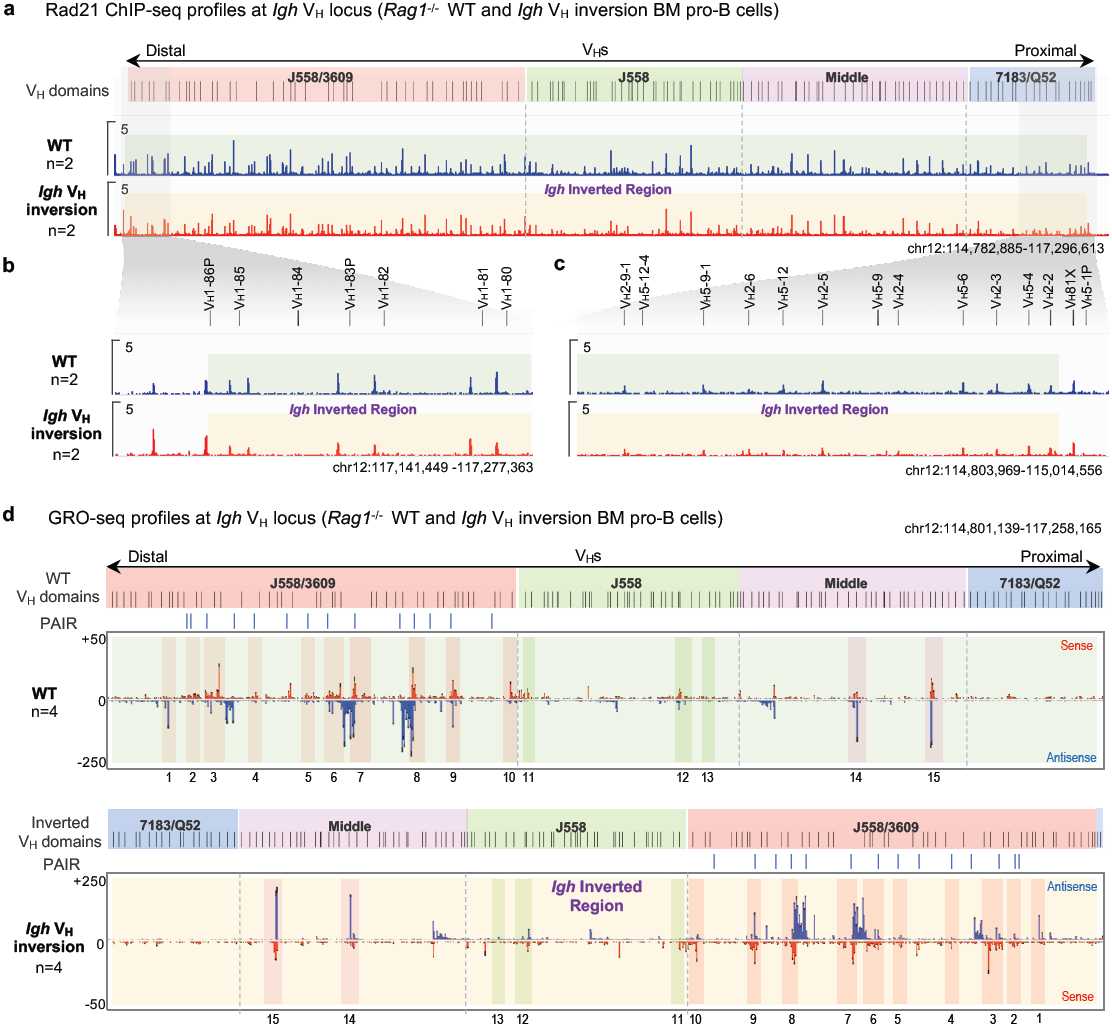
Rad21-binding and transcription across *Igh* locus in WT and *Igh* V_H_ locus inversion cultured *Rag1*^*-/-*^ BM pro-B cells. **a**, Representative Rad21 ChIP-seq profiles across the 4 V_H_ domains as indicated in WT (upper, blue) and *Igh* V_H_ locus inversion (bottom, red) primary pro-B cells. **b, c**, Rad21 ChIP-seq profiles zoomed in the distal *Igh* V_H_ region (**b**) and proximal *Igh* V_H_ region (**c**). **d**, Average signal counts ± s.e.m. of GRO-seq profiles across the 4 V_H_ domains in WT (upper) and *Igh* V_H_ locus inversion (bottom) BM pro-B cells. The WT and inverted V_H_ locus/domains with PAIR elements are diagrammed at the top of each panel. Both the sense and antisense transcription are relative to the entire *Igh* V_H_ locus upstream V_H_81X with or without inversion and indicated respectively. For comparison, 15 representative major interaction peaks/clusters as shown in Fig.1c were indicated with colour shades and numbers. The background of *Igh* V_H_ inverted region is highlighted in transparent green (WT) and transparent gold (*Igh* V_H_ inversion), respectively. In all panels, “n” indicates biological repeats. See Methods for more details.

**Extended Data Figure 4.**
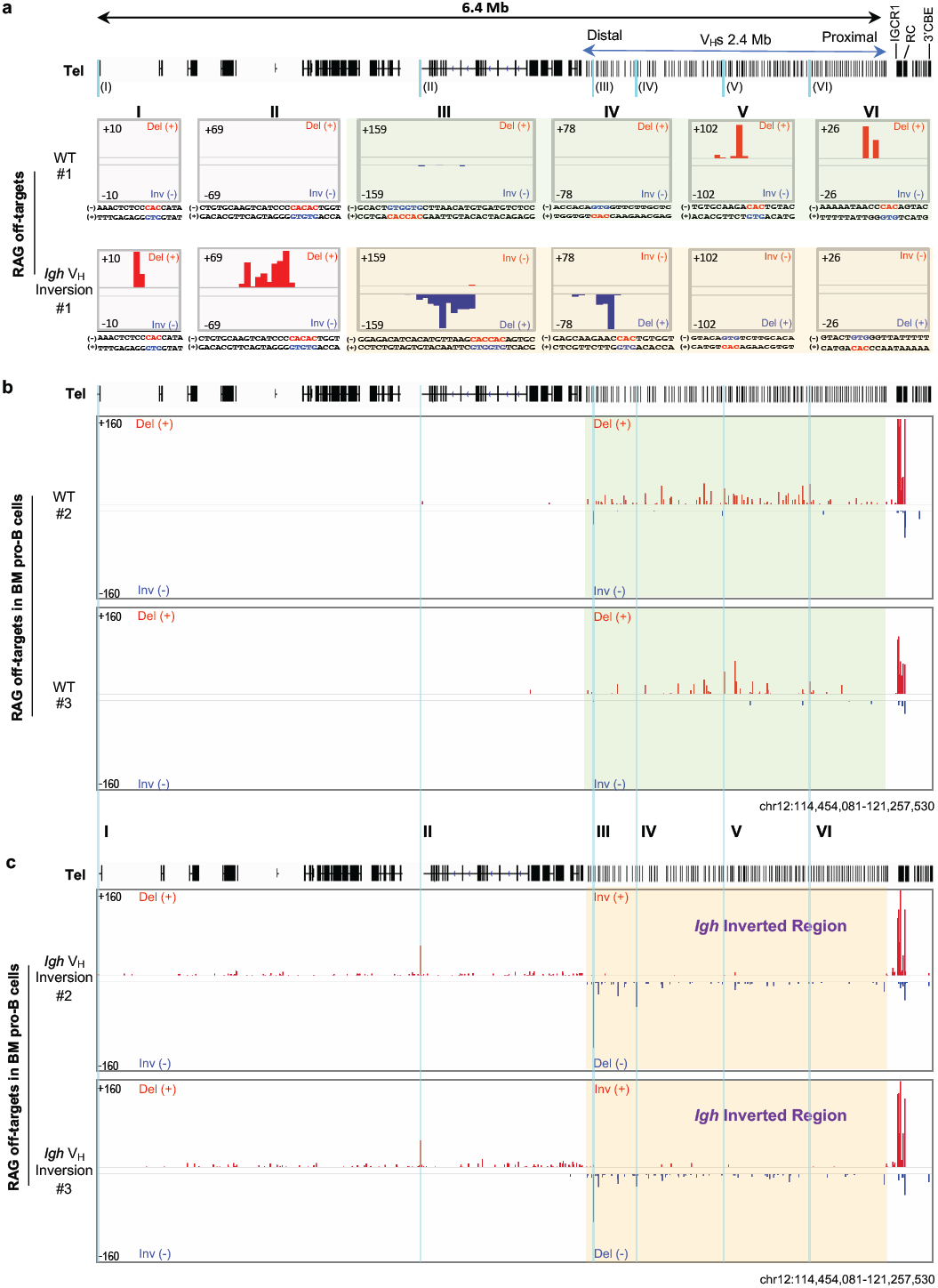
RAG off-target profiles in WT and *Igh* V_H_ locus inversion primary BM pro-B cells. **a**, 6 examples (labeled in Fig. 2b, I-VI) of RAG off-target peaks in WT (repeat#1, middle) and *Igh* V_H_ locus inversion (repeat#1, bottom) BM pro-B cells plotted at single-base-pair resolution. Top panels are WT 2.4 Mb *Igh* locus and upstream 4 Mb region track. **b, c**, RAG off-target junction profiles at *Igh* locus and upstream 4 Mb region in WT (repeat#2-3, middle) and *Igh* V_H_ locus inversion (repeat#2-3, bottom) BM pro-B cells isolated from individual mice. The same regions of RAG off-target peaks (shown in **a**) are highlighted with sky blue lines. #1-3, number of individual WT or *Igh* V_H_ inversion mice. The background of *Igh* V_H_ inverted region is highlighted in transparent green (WT) and transparent gold (*Igh* V_H_ inversion), respectively. Del (+), deletional junction. Inv (-), inversional junction.

**Extended Data Figure 5.**
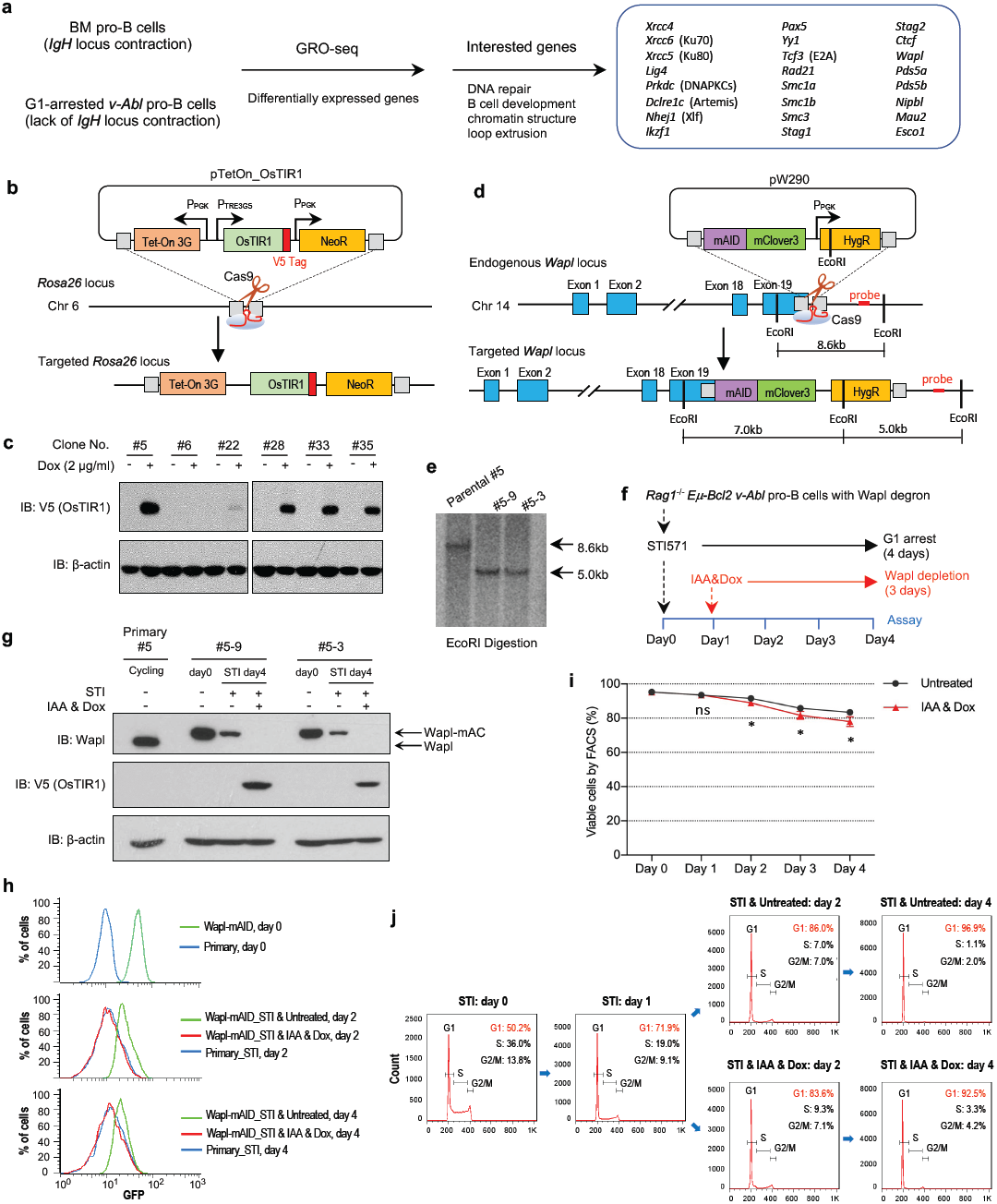
Generation and characterization of Wapl-degron *v-Abl* pro-B lines. **a**, Strategy for detecting differentially expressed genes between BM pro-B cells and G1-arrested *v-Abl* pro-B cells by GRO-seq. 24 interested genes are listed. **b**, Schematic of the targeting strategy for introducing Tet-On OsTIR1 expression cassette at the mouse *Rosa26* locus. Positions of homology arms (gray box) are indicated. Cells were transfected with pTet-On_OsTIR1_V5 and pX330_Rosa26 to target *Rosa26* locus. The clones were screened by genomic PCR. See Methods for more details. **c**, Immunoblotting to detect OsTIR1 expression with the induction of Doxycycline (Dox). The indicated clones were grown in the presence of 2 mg/ml Dox for 24 hr before immunoblotting with the anti-V5 antibody. Primary #5 clone was picked for further Wapl-degron targeting. β-actin was a loading control. **d**, Strategy to generate Wapl-degron *v-Abl* pro-B cell lines. Cells were transfected with pW290 and pX330_Wapl-AID to introduce in-frame mAID-Clover sequences into the last codon of mouse both endogenous *Wapl* alleles. Positions of homology arms (gray box), Cas9/sgRNAs and Southern blot probes are indicated. **e**, Southern blot confirmation of two correctly targeted clones (#5-3 and #5-9) with Wapl-mAID on both alleles. **f**, Diagram of the experimental strategy to specifically deplete mAID-tagged Wapl protein in G1-arrested *v-Abl* cells. **g**, Immunoblotting to detect Wapl and Wapl-mAID-Clover (Wapl-mAC) protein. The indicated clones (#5-3 and #5-9) were grown without or with Wapl depletion at indicated time points before immunoblotting. The specific western blot bands of WT Wapl and Wapl-mAC were labeled. OsTIR1 was detected by anti-V5 antibody. Primary #5 clone was used for the WT Wapl control and β-actin was a loading control. **h**, Representative flow-cytometry plots showing the percentage of Clover-positive Wapl-degron *v-Abl* cells that are without (Untreated) or with (IAA&Dox) Wapl depletion at different time points (day 0, top; day 2, middle; day 4, bottom). Primary line #5 was processed as a Clover-negative control. **i**, Time-course cell viability assay for G1-arrested *v-Abl* pro-B cells without or with Wapl depletion at different time points. Average percentage ± s.d. of viable cells for each timepoint and for each condition was calculated 4 independent experiments. **j**, Representative flow-cytometry plots of propidium iodide (PI) stained G1-arrested *v-Abl* pro-B cells without or with Wapl depletion at indicated time points. Percentages in the top-right corner represent the percentage of cells at G1, S and G2/M stage. Plasmids, sgRNAs and oligos used are listed in Supplementary Table 4. *p* values were calculated using unpaired two-tailed t-test. See Methods for more details.

**Extended Data Figure 6.**
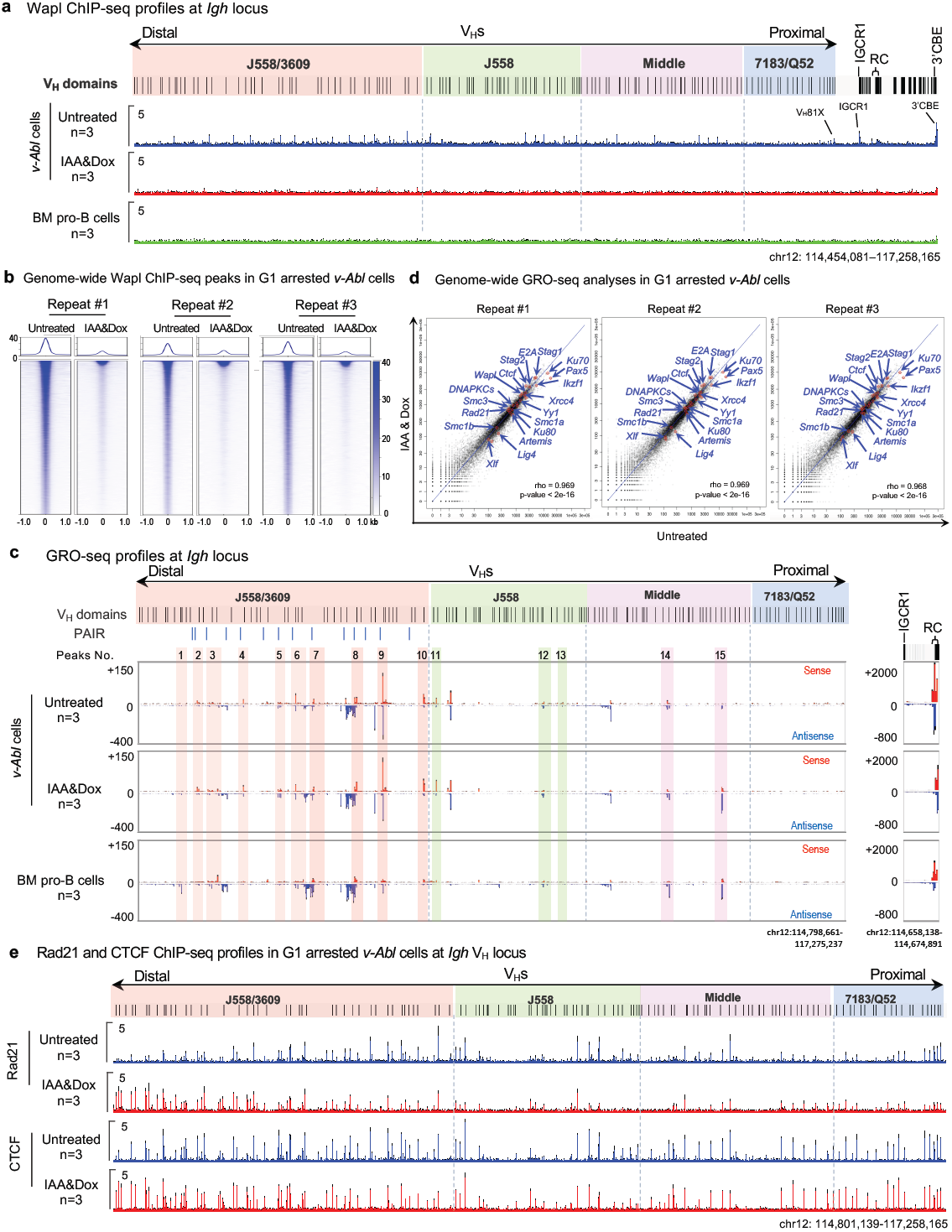
Characterization of Wapl/Rad21/CTCF-binding and genome-wide gene transcription in G1-arrested Wapl-degron *v-Abl* pro-B cells. **a**, Average signal counts ± s.e.m. of Wapl ChIP-seq across the *Igh* locus are plotted as indicated in *Rag1*^*-/-*^ G1-arrested *v-Abl* pro-B cells without (Untreated, blue, n=3) or with (IAA&Dox, red, n=3) Wapl depletion and cultured *Rag1*^*-/-*^ BM pro-B cells (green, n=3). **b**, Three independent repeats of Wapl ChIP-seq signal within ±1.0 kb region across all peaks genome-wide called in G1-arrested *Rag1*^*-/-*^ *v-Abl* pro-B cells without (Untreated) or with (IAA&Dox) Wapl depletion. Top: Average enrichment. IAA&Dox treatment leads to a depletion of chromatin-bound Wapl genome-wide. See Methods for details. **c**, Average signal counts ± s.e.m. of GRO-seq across the 4 V_H_ domains (left) and RC region (right) are plotted as indicated in G1-arrested *Rag1*^*-/-*^ *v-Abl* pro-B cells without (Untreated, upper, n=3) or with (IAA&Dox, middle, n=3) Wapl depletion and cultured *Rag1*^*-/-*^ BM pro-B cells (bottom, n=4). For comparison, 15 representative major interaction peaks/clusters as Fig.1c were indicated with colour shades and numbers. PAIR elements are indicated as blue Bars. **d**, Scatter plots of transcriptome-wide GRO-seq counts from three independent repeats in G1-arrested *Rag1*^*-/-*^ *v-Abl* pro-B cells without (Untreated, *x* axis) and with (IAA&Dox, *y* axis) Wapl depletion. Representative known requisite genes for V(D)J recombination and chromatin interaction are highlighted by red circles and blue arrows in each of the three scatter plots. See Methods for more details. **e**, Average signal counts ± s.e.m. of Rad21 (upper) and CTCF (bottom) ChIP-seq across the four 4 V_H_ domains are plotted as indicated in G1-arrested *Rag1*^*-/-*^ *v-Abl* pro-B cells without (Untreated, blue, n=3) or with (IAA&Dox, red, n=3) Wapl depletion. See Methods for more details.

**Extended Data Figure 7.**
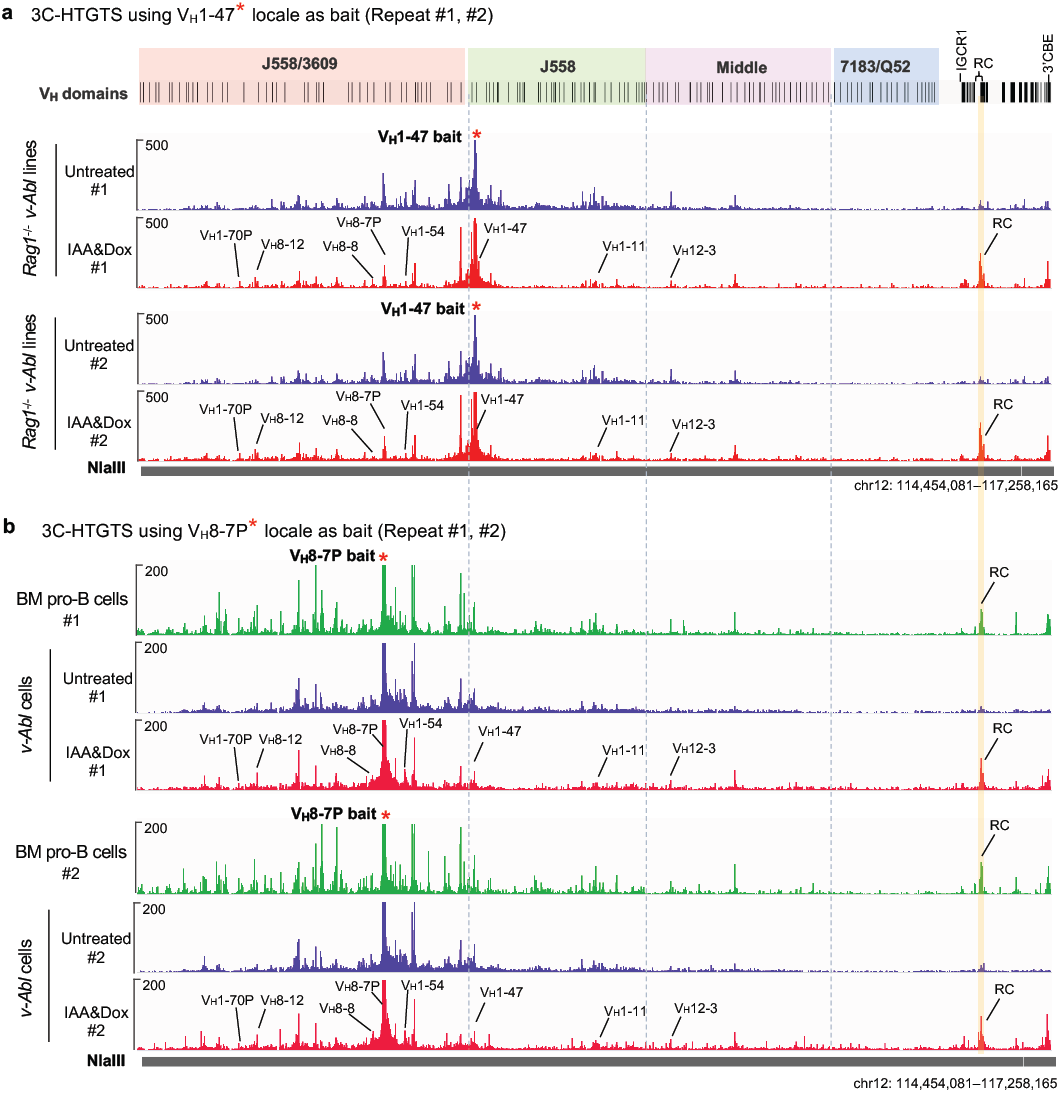
Wapl depletion results in increased chromatin interactions of the RC with distal V_H_s in G1-arrested Wapl-degron *v-Abl* pro-B cells. **a**, 3C-HTGTS chromatin interaction profiles of distal V_H_1-47 bait across the entire *Igh* locus in G1-arrested *Rag1*^*-/-*^ *v-Abl* pro-B cells without (Untreated, blue, n=2) or with Wapl depletion (IAA&Dox, red, n=2). **b**, 3C-HTGTS chromatin interaction profiles of distal V_H_8-7P bait across the entire *Igh* locus in WT *Rag1*^*-/-*^ mouse BM pro-B cells (BM, green, 2 repeats) and G1-arrested *Rag1*^*-/-*^ *v-Abl* pro-B cells without (Untreated, blue, n=2) or with Wapl depletion (IAA&Dox, red, n=2). Related very low-level peaks of RC interactions in untreated cell suggest low-level *Igh* locus contraction in untreated cells. “n” indicates experimental repeats. 2 biological repeats for BM pro-B cells. See Methods for more details.

**Extended Data Figure 8.**
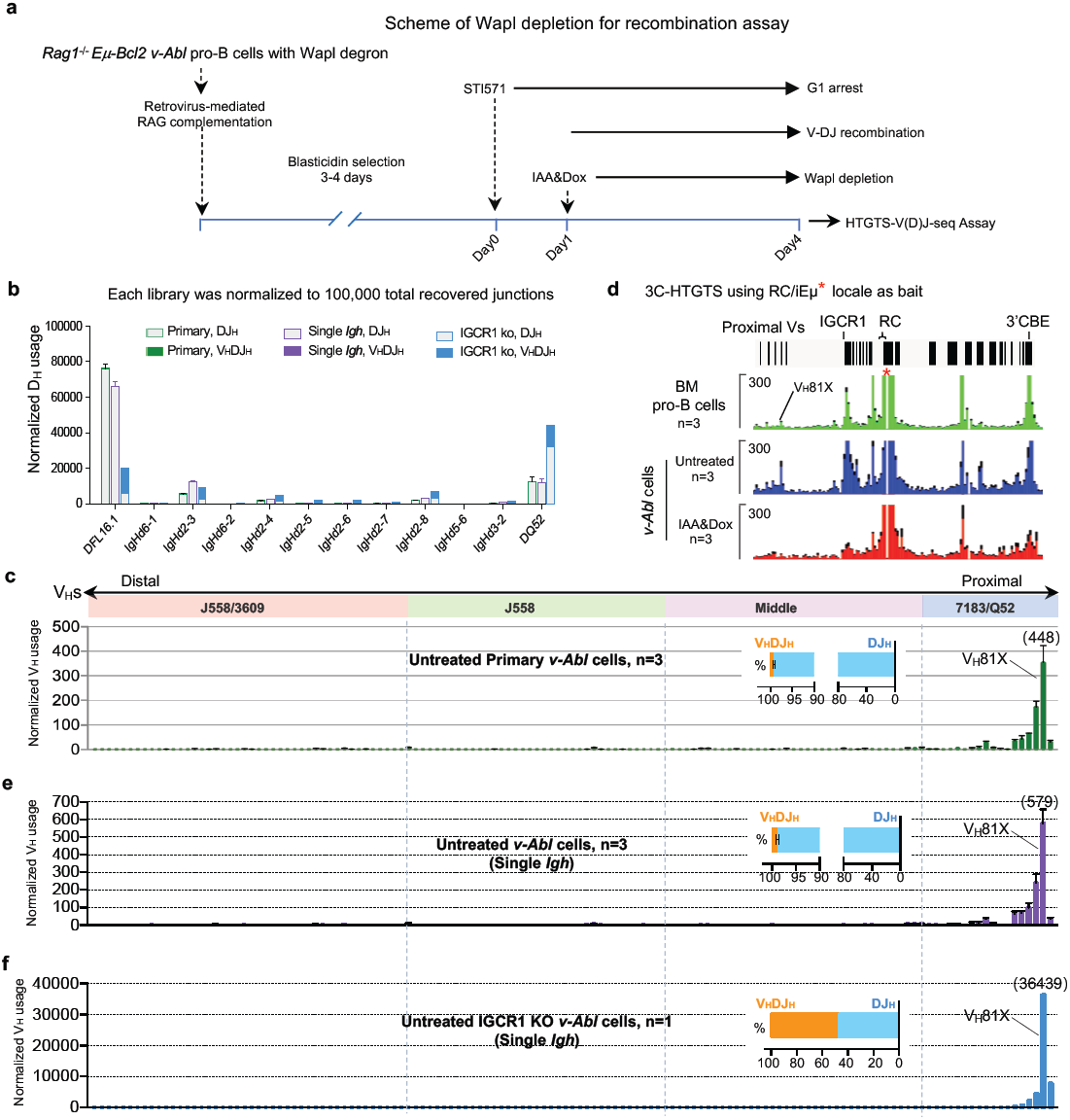
HTGTS-V(D)J-seq assay in RAG1-complemented, G1-arrested Wapl-degron *v-Abl* pro-B cells. **a**, Diagram of the experimental strategy with retrovirus-mediated RAG complementation in Wapl-degron *v-Abl* cells for HTGTS-V(D)J-seq assay. See Methods for more details. **b**, Average utilization frequencies ± s.d. of all D segments from DJ_H_ and V_H_DJ_H_ joins in RAG-complemented, G1-arrested untreated primary *v-Abl* cells (primary, green, n=3), untreated single *Igh* Wapl-degron *v-Abl* pro-B cells (single *Igh*, purple, n=3) and untreated single *Igh* with IGCR1 KO Wapl-degron *v-Abl* pro-B cells (IGCR1 KO, blue, n=1). **c**, Average utilization frequencies ± s.d. of all V_H_ segments in RAG1-complemented, G1-arrested primary *v-Abl* pro-B cells (n=3). Average percentage ± s.d. of V_H_DJ_H_ and DJ_H_ rearrangements are indicated**. d**, Average 3C-HTGTS signal counts ± s.e.m. baiting from RC/iEμ across the locus between 3’CBE and several proximal Vs in cultured *Rag1*^*-/-*^ BM pro-B cells (top, green, n=3), G1-arrested *Rag1*^*-/-*^ *v-Abl* pro-B cells without (middle, blue, n=3) or with Wapl depletion (bottom, red, n=3). Related very low-level peaks of RC interactions with IGCR1 in IAA/Dox treated cells suggest IGCR1 was neutralized in Wapl depleted *v-Abl* cells. **e**, Average utilization frequencies ± s.d. of all V_H_ segments in RAG1-complemented, G1-arrested untreated single *Igh* Wapl-degron *v-Abl* pro-B cells (n=3). Average percentage ± s.d. of V_H_DJ_H_ and DJ_H_ rearrangements are indicated**. f**, Representative utilization frequencies of all V_H_ segments in RAG1-complemented, G1-arrested untreated single *Igh* with IGCR1 KO Wapl-degron *v-Abl* pro-B cells (n=1). Representative percentage of V_H_DJ_H_ and DJ_H_ rearrangements are indicated. All V_H_ segments divided into four domains from most proximal to distal. *p* values were calculated using unpaired two-tailed t-test. See Supplementary Table 2, 3 and Methods for more details.

**Extended Data Figure 9.**
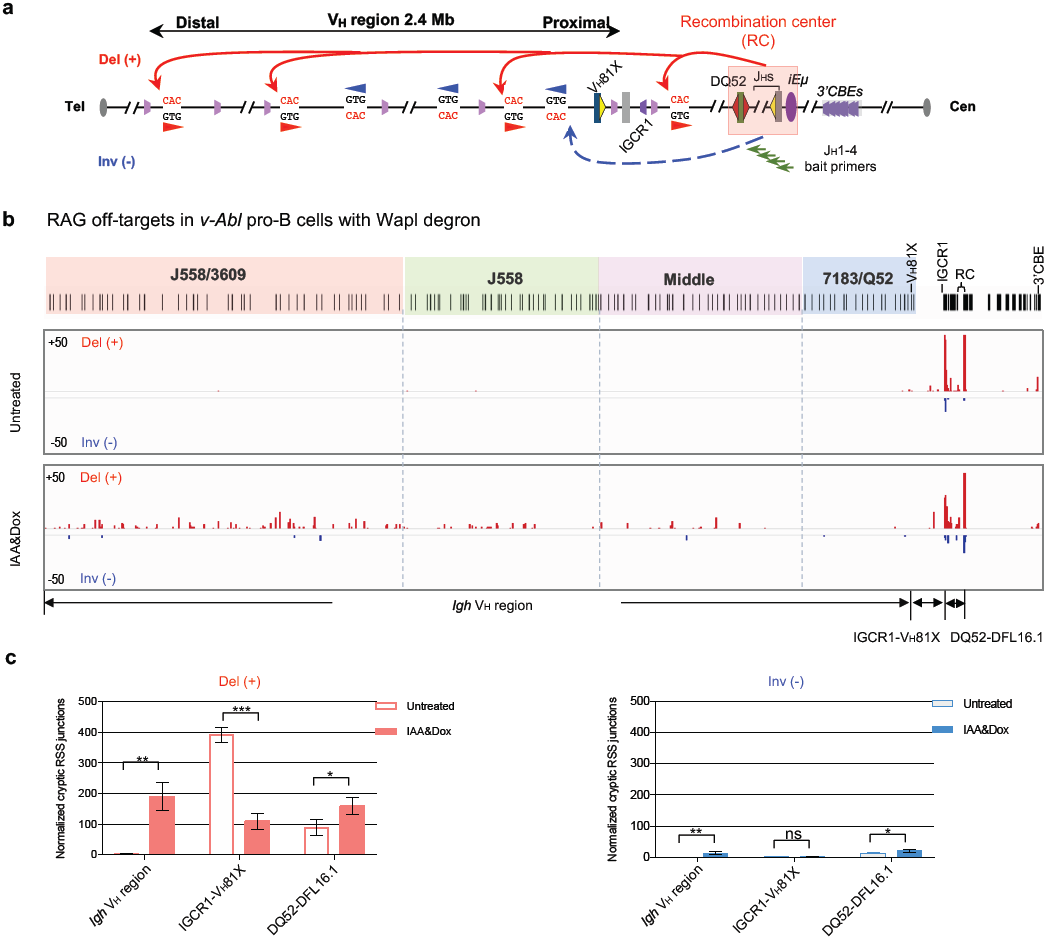
Substantial Wapl depletion promotes robust RAG utilization of cryptic RSSs across the V_H_ locus in G1-arrested Wapl-degron *v-Abl* pro-B cells. **a**, Illustration of possible joining outcomes between *bona fide* RSSs from J_H_1-4 coding end baits to cryptic RSSs mostly represented by CAC motifs across *Igh* locus, details as shown in Fig2a. **b**, Pooled HTGTS junction profiles across highlighted V_H_ domains for deletional and inversional joining in RAG1-complemented, G1-arrested *v-Abl* pro-B cells without (Untreated, upper, pooled n=3) or with Wapl depletion (IAA&Dox, bottom, pooled n=3). For presentation, all libraries from untreated or IAA&Dox treated cells were pooled and normalized. See Methods for more details. The vast majority of deletional joining events across the V_H_ locus are reproducible among replicates and involve, at a minimum the CAC of an RSS. Almost all of the handful of the very low-level inversional joining events interspersed across V_H_ locus are not reproducible among replicates consistent with being background events. **c**, Average frequencies ± s.d. of deletional (left) and inversional (right) joining events within indicated regions in RAG1-complemented, G1-arrested *v-Abl* pro-B cells without (Untreated, n=3) or with Wapl depletion (IAA&Dox, n=3). Each replicate was normalized to 500 off-target junctions for statistical analysis. Three indicated regions (IgH V_H_ region, IGCR1-V_H_81X and DQ52-DFL16.1) were labeled (**b**), respectively. Del (+), deletional junction. Inv (-), inversional junction. *p* values were calculated using unpaired two-tailed t-test. See Methods for more details.

## Additional Information for Supplementary Tables and Data

**Supplementary Table 1 |** Relative utilization of *Igh* V and D segments in WT and *Igh* V_H_ locus inversion primary BM pro-B cells (see text).

**Supplementary Table 2 |** Relative utilization of *Igh* V and D segments in RAG1-complemented, G1-arrested, primary *v-Abl* pro-B cells and Wapl-degron *v-Abl* pro-B cells without or with Wapl depletion (see text).

**Supplementary Table 3 |** Relative utilization of *Igh* V and D segments in IGCR1-intact or IGCR1-deleted RAG1-complemented G1-arrested untreated single *Igh* Wapl-degron *v-Abl* pro-B cells (see text).

**Supplementary Table 4 |** Oligos and plasmids used in this study.

**Supplementary Data 1 |** 3C-HTGTS, GRO-seq, ChIP-seq, CBE motif sites and PAIR elements for 15 representative peaks/clusters in V_H_s domains in cultured *Rag1*^*-/-*^ BM pro-B cells (see text).

**Supplementary Data 2 |** V_H_ usage, GRO-seq, 3C-HTGTS, ChIP-seq, CBE motifs sites and PAIR elements for all V_H_s in G1-arrested Wapl-degron *v-Abl* pro-B cells (see text).

## METHODS

### Experimental procedures

No statistical methods were used to predetermine sample size. Experiments were not randomized, and the investigators were not blinded to allocation during experiments and outcome assessment.

### Mice

Wild-type (WT) C57BL/6 mice were purchased from Charles River Laboratories International. *Rag1*^-/-^ mice (stock No.002216: B6.129S7-*Rag1*^tm1Mom^/J) in C57BL/6 background were purchased from the Jackson Laboratory and maintained in the lab. All animal experiments were performed under protocols approved by the Institutional Animal Care and Use Committee of Boston Children’s Hospital.

### Generation and characterization of *Igh* V_H_ locus inversion mouse model

The CRISPR/Cas9-mediated entire *Igh* V_H_ locus inversion upstream V_H_81X modifications were carried out on the C57BL/6 *Igh* allele in EF1 ES cell line, which was derived from a F1 hybrid mouse (129/Sv:C57BL/6). The ES targeting was performed as previously described^36^. The cells were cultured at 37 °C with 5% CO_2_ in DMEM medium supplemented with 15% fetal bovine serum, 20 mM HEPES, 1× MEM nonessential amino acids, 2 mM Glutamine, 100 units of Penicillin/Streptomycin, 100 μM β-mercaptoethanol, 500 units/ml Leukemia Inhibitory Factor (LIF). The ES cells were grown on a monolayer of mouse embryonic fibroblasts (MEF) that have been mitotically inactivated with γ-irradiation. sgRNAs targeting the upstream of *V*_*H*_*1-86P* (Cut1) and upstream of *V*_*H*_*81-X* (downstream of *V*_*H*_*2-2*) (Cut2) were cloned by annealing pairs of oligos into pX459 with a puromycin selection marker (Addgene, #62988) following the standard protocol. For transfection, Cas9/sgRNAs (Cut1 and Cut2, each 2.5 μg) were nucleofected via Lonza 4D Nucleofector into 2.5×10^6^ EF1 cells. The electroporated ES cells were grown on a monolayer of MEF cells with puromycin resistance (Stemcell, #00325) for 1 day. Then the ES cells were selected with 2 μg/ml puromycin (Gibco, #A1113802) for another day. After selection, the ES cells were cultured with normal medium for 5-6 days, colonies were picked into 24 well plates that have been coated with MEF. After 3 days, half of the colonies were frozen at -80°C and the other half were expanded for DNA isolation. The genomic DNA was analyzed with screening-PCR to identify clones in which the desired genetic modifications have taken place. After the primary screen, the upstream and downstream of modified region of the potential positive clones was amplified by confirming-PCR and sequenced. After the *Igh* V_H_ inversion modifications were confirmed on the C57BL/6 *Igh* allele by single nucleotide polymorphisms (SNP), each positive clone was further validated with a normal karyotype by metaphase analysis. All the cell lines were confirmed to be mycoplasma free. The ES clone with the *Igh* V_H_ inversion modifications was injected into *Rag2*^*-/-*^ blastocysts to generate RDBC chimeras^37^. The chimeric mice were bred with C57BL/6 mice for germline transmission of the targeted mutations. Heterozygous *Igh* V_H_ inversion mice were bred to homozygosity (*Igh* V_H_ inversion mouse model). For further analysis, this mouse model was bred to *Rag1*^-/-^ homozygosity in C57BL/6 background. Sequences of all sgRNAs and oligos used are listed in Supplementary Table 4.

### Bone marrow pro-B cells purification

For RAG on-target and off-target analysis, single cell suspensions were derived from bone marrows of 4-6-week-old C57BL/6 WT and *Igh* V_H_ locus inversion mice and incubated in Red Blood Cell Lysing Buffer (Sigma-Aldrich, #R7757) to deplete the erythrocytes. B220^+^CD43^hi+^IgM^-^ pro-B cells were isolated by staining with anti-B220-APC (eBioscience, #17-0452-83), anti-CD43-PE (BD PharMingen, #553271), and anti-IgM-FITC (eBioscience, #11-5790-81) antibodies for 30 minutes at 4 °C and then purified by FACS, the purified primary pro-B cells were subjected to HTGTS-V(D)J-seq.

For 3C-HTGTS, ChIP-seq and GRO-seq, B220-positive WT and *Igh* V_H_ inversion pro-B cells were separately purified via anti-B220 MicroBeads (Miltenyi, #130-049-501) from individual 4-6-week-old *Rag1* deficient mice (WT and *Igh* V_H_ locus inversion) and were cultured in opti-MEM medium containing 10% (v/v) FBS plus 10 ng/mL IL-7 and 2 ng/mL SCF for 4-5 days as previously described^38^. Purified BM pro-B cells from different mice were separately cultured in different Petri dishes.WT and *Igh* V_H_ inversion mice and short-term cultured BM-pro-B cells were double-checked and confirmed by PCR, respectively, prior to various assays as described below. PCR primers are listed in Supplementary Table 4.

### Generation of doxycycline inducible Wapl-degron *v-Abl* pro-B lines

The targeting plasmid (pTet-On_OsTIR1_V5) for introducing the doxycycline inducible OsTIR1-V5 expression cassette into endogenous *Rosa26* locus was constructed by modifying a published pEN113 plasmid (Plasmid #86233, Addgene). The donor plasmid (pW290) used to target the endogenous mouse *Wapl* locus was constructed by modifying a published pMK290 plasmid (Plasmid #86230, Addgene). Briefly, the doxycycline inducible Tet-On-3G_hPGK promoter sequence was PCR amplified from the plasmid pMK243 (Addgene, #72835) with 5’ SalI and 3’ MluI digestion sites. Then the pTet-On_OsTIR1_V5 targeting vector was generated by cutting out the enhancer and promoter sequence of OsTIR1_V5 expression cassette from the pEN113 plasmid via SalI and MluI digestion, and replacing the amplified Tet-On-3G_hPGK promoter sequence into pEN113 vector via the same restriction sites (Extended Data Fig.5b). To construct donor pW290 vector, 5’ and 3’ homology arms (247 bp each) flanking the BamHI sequence were synthesized via gBlock by IDT (Integrated DNA Technologies) and cloned into pGEM-T easy vector (Promega, #A1360) to generate the pGEM-Wapl plasmid. The mAID-mClover cassette containing a hygromycin selection marker was excised from pMK290 by BamHI and cloned at the BamHI site of the pGEM-Wapl plasmid, between the homology arms, to generate the pW290 vector (Extended Data Fig. 5d). To modify the mouse *Rosa26* and *Wapl* gene, sgRNAs targeting the endogenous *Rosa26* locus or 3’ end sequence of the *Wapl* gene were cloned by annealing pairs of oligos into pX330 (Addgene, #42230) to construct the pX330_Rosa26 and pX330_Wapl-mAID, respectively. Sequences of all sgRNAs and oligos used are listed in Supplementary Table 4.

The *Rag1*^-/-^; *Eμ-Bcl2*^*+*^ *v-Abl* pro-B cells were co-transfected with pTet-On_OsTIR1_V5 and pX330_Rosa26 via Lonza 4D Nucleofector as shown before^9,10^, selected with 0.4 mg/ml of G418 for 5-6 days, and subcloned by dilution. Candidate clones with desired gene modifications were screened by PCR. The expression of OsTIR1 was confirmed by Western blot (Extended Data Fig. 5c). Similarly, the resultant clones were further co-transfected with donor pW290 with corresponding Cas9/sgRNA pX330_Wapl-mAID via nucleofection, selected with 800 μg/ml hygromycin B (Millipore Sigma, #10843555001) for 3 days, and subcloned by dilution. Candidate clones with desired gene modifications were screened by PCR and confirmed by Southern blot in Extended Data Fig. 5d-e, and the expression of mAID-tagged Wapl protein was confirmed by Western blot and FACS as outlined in Extended Data Fig. 5f-j. The resulting lines were referred to as the Wapl-degron v*-Abl* pro-B lines and used in this study.

### Treatment of Wapl-degron *v-Abl* pro-B lines with IAA&Dox

To specifically deplete mAID-tagged Wapl protein in the G1 stage, Wapl-degron *v-Abl* pro-B lines were treated with 150 μM Indole-3-acetic acid (auxin analog, IAA, Sigma-Aldrich, #I3750-25G-A) and 2 mg/mL doxycycline (Dox, Sigma-Aldrich, #D3447) from day1 to day4 during 3 μM STI-571 (STI) induced four-day G1 arrest experimental period (day0-4) (Extended Data Figure. 5f). All the chemicals were dissolved in DMSO to prepare the stock. DMSO solvent was applied as the mock to untreated cells. Both untreated and IAA&Dox-treated cells were then collected and examined by FACS for protein depletion confirmation (Extended Data Figure. 5h) prior to various assays as described below.

### Western Blot

The western blot experiment was performed according to published protocol with modification^39^. Before harvesting, the primary line with Tet-on OsTIR1-V5 tag were treated with/without doxycycline for 24 hr and the Wapl-degron *v-Abl* cells were treated with/without IAA&Dox as described in Extended Data Fig. 5f. 10 million cells were harvested and lysed in EBC buffer (50 mM Tris-HCl pH 7.5, 120 mM NaCl, 0.5% NP40) with protease inhibitors (Complete Mini, Roche) for 30 min on ice, followed by pulse sonication for 5 s with low energy input. The lysates were then resolved by SDS-PAGE and immunoblotted with indicated antibodies (Wapl and V5-tag antibodies were purchased from Thermo Fisher Scientific #PA5-38024 and #R960-25, and β-actin antibody was purchased from Cell Signaling Technology #3700).

### Cell viability assay

For time-course cell viability assay, Wapl-degron *v-Abl* cells were treated with IAA&Dox from day1 to day4 during the STI-571 induced four-day G1 arrest experimental period (day0-4). The viability of Wapl-degron *v-Abl* pro-B cells with or without Wapl depletion at different time points (day 0 to day 4) was determined by the percentage of viable lymphocyte population gated by FACS side (SSC) and forward (FSC) scatters out of the total cells. Average percentage ± s.d. of viable cells for each timepoint and each condition was calculated 4 independent experiments.

### GFP quantification and cell cycle stage analysis

For GFP (Clover) signaling quantification of Wapl depletion, both the untreated and IAA&Dox-treated Wapl-degron *v-Abl* cells at different time points (day0, day2 and day4) were directly collected and examined by FACS. Primary *v-Abl* line #5 was processed as a GFP negative control.

For cell cycle analysis, Wapl-degron *v-Abl* pro-B cells without or with Wapl depletion at different time points (day0, day1, day2 and day4) were pelleted at 500 g for 5 min at 4 °C, washed with ice-cold PBS, pelleted again, then fixed in 66% ethanol on ice: gently resuspend the cell pellet in 400 μL ice-cold 1х PBS. Slowly add 800 μL ice-cold 100% ethanol and mix well. Store at 4 °C for at least 2 hours. Gently resuspended the fixed cell pellet in 300 μL 50 μg/mL propidium iodide (PI) and 0.2 mg/mL RNase in PBS. After 30 min incubation in the dark at 37 °C, cells were transferred on ice, passed through an appropriate filter to remove cell aggregates and then used directly for flow cytometry.

### RAG complementation

The *Rag1*-expressing vector pMSCV-RAG1-IRES-Bsr and *Rag2*-expressing vector pMSCV-FLAG-RAG2-GFP were kindly provided by Yu Zhang and Xuefei Zhang. RAG was reconstituted in RAG1-deficient *v-Abl* cells via retroviral infection of cells with the pMSCV-RAG1-IRES-Bsr and pMSCV-FLAG-RAG2-GFP vectors followed by 3-4 days of Blasticidin (ThermoFisher, #R21001) selection to enrich for cells with virus integration.

### HTGTS-V(D)J-seq

For *Igh* V(D)J recombination analyses, we purified BM pro-B cells from 4-6-week-old WT C57BL/6 mice and entire *Igh* V_H_ locus inversion mice as described above and introduced RAG into *Rag1*^*-/-*^ Wapl-degron *v-Abl* pro-B cells as well as their derivatives via the approach described previously. HTGTS-V(D)J-seq libraries were prepared as previously described^1,10,17,18^. Briefly, 1 μg of gDNA from sorted mouse BM pro-B cells or 8 μg of gDNA from G1-arrested RAG-complemented Wapl-degron *v-Abl* cells with or without Wapl depletion was sonicated and subjected to LAM-PCR using biotinylated J_H_1-4 bait primers. Single-stranded LAM-PCR products were purified using Dynabeads MyONE C1 streptavidin beads (Life Technologies, #65002) and ligated to bridge adaptors. Adaptor-ligated products were amplified by nested PCR with indexed J_H_1-4 primers and the primer annealed to the adaptor. The PCR products were further tagged with Illumina sequencing adaptor sequences, size-selected via gel extraction and loaded onto an Illumina NextSeq550 using paired-end 150-bp sequencing kit. Primer sequences are listed in Supplementary Table 4.

### HTGTS-V(D)J-seq data processing and analyses

HTGTS-V(D)J-seq libraries were processed via the pipeline described previously^18^. The data were aligned to mm9 genome and analyzed with all duplicate junctions included in the analyses as previously described. In the *Igh* repertoire libraries generated from C57BL/6 WT BM pro-B cells with J_H_1-4 coding end baits^17,40^, we detected in V_H_DJ_H_ exons 109 functional V_H_s, as well as 23 pseudo V_H_s (Supplementary Table 1) (average rearrangement >1).For comparisons, utilization data of V_H_ and D segments in WT or *Igh* V_H_ inverted BM pro-B cells was normalized to 118,475 total recovered junctions (Fig. 1b; Supplementary Table 1). Utilization data of V_H_ and D segments in G1-arrested Wapl-degron *v-Abl* cells was normalized to 100,000 total recovered junctions (Fig. 3e; Extended Data Fig. 8b, c; Supplementary Table 2). Utilization data of V_H_ and D segments in G1-arrested single *Igh* Wapl-degron *v-Abl* cells with or without IGCR1 was normalized to 100,000 total recovered junctions (Extended Data Fig. 8b, e, f; Supplementary Table 3). For the RAG off-targets in WT or *Igh* V_H_ inverted BM pro-B cells, each library was isolated from individual independent J_H_1-4 primer libraries and was normalized to 402,241 isolated junctions (Fig. 2b; Extended Data Fig. 4). For comparison, each of the biological library replicates in BM pro-B cells was normalized to 2979 off-target junctions in the indicated region (chr12:114,666,712-121,257,530) for statistical analysis (Fig. 2c). For the RAG off-targets in Wapl-degron *v-Abl* cells and derived *Igh* V_H_ inverted lines, each library was pooled from 3 independent J_H_1-4 primer libraries and was normalized to 432,676 isolated junctions without repetitive sequences (Extended Data Fig. 9b). For comparison, each of the biological library replicates in *v-Abl* lines was normalized to 500 off-target junctions in the indicated region (DQ52-entire *Igh* V_H_) for statistical analysis (Extended Data Fig. 9c). Junctions are denoted as in ‘‘+’’ orientation if prey sequence reads in centromere-to-telomere direction and in ‘‘-’’ orientation if prey sequence reads in telomere-to-centromere direction (Fig. 2; Extended Data Fig. 9). Data reproducibility was controlled by performing at least three independent experiments and showed as mean ± s.d. for visualization.

### ChIP-Seq library preparation and analyses

ChIP-seq libraries were prepared with short-term cultured *Rag1*^*-/-*^ BM pro-B cells or G1-arrested *Rag1*^*-/-*^ *v-Abl* pro-B cells (with or without Wapl depletion) and performed based on previously description with modification^2^. In brief, 20 million cells were crosslinked in 37 °C prewarmed culture medium with 1% formaldehyde for 10 min at 37 °C and quenched with glycine at a final concentration of 125 mM. Cells were then treated with cell lysis buffer (5 mM PIPES pH 8.0, 85 mM KCl, 0.5% NP40) for 10 min on ice, followed by treatment with nuclear lysis buffer (50 mM Tris-HCl pH 8.1, 10 mM EDTA, 1% SDS) for 10 min at room temperature. Chromatin was subjected to sonication with Diagenode Bioruptor at 4 °C to achieve an average size of 200-300 bp (30s on, 30 s off, 20 cycles with high energy input). Chromatin was then precleared with Dynabeads Protein A (Thermo Fisher Scientific, 10002D) at 4 °C for 2 h. 50 mL lysates was kept as input and the rest were incubated with 5 μg RAD21 antibody (Abcam, #ab992) or 10 μg CTCF antibody (Millipore, #07-729) or 5 μg Wapl antibody (Thermo Fisher Scientific, #PA5-38024) overnight at 4 °C. Immunoprecipitation (IP) samples were then captured by Dynabeads Protein A at 4 °C for at least 2 h followed by bead washing and elution. IP and input DNA were de-crosslinked at 65 °C overnight and purified via Qiagen PCR purification columns. Purified DNA was subjected to ChIP-Seq library preparation with NEBNext Ultra II DNA Library Prep Kit for Illuumina (NEB, #E7645) and sequenced by paired-end 75-bp sequencing on Next-Seq550. For comparison of all the ChIP-seq results in this study, we used bowtie2 to align ChIP-seq reads to mm9 genome and run MACS2 callpeak with parameters ‘-t IP.bam -c input.bam –nomodel –keep-dup all -extsize 51 -nolambda -B -SPMR -g mm -broad’ to generate bigwig graph normalized to 10 million reads. For comparison of ChIP-seq results between WT and *Igh* inversion BM pro-B cells, Rad21 ChIP-seq profiles across the 4 V_H_ domains were potted in the normal orientation (Extended Data Fig. 3a-c). Indicated replicates were combined as mean ± s.d. for visualization.

### GRO-seq and data analyses

GRO-seq libraries were prepared as described previously^1,41^ on short-term cultured *Rag1*^*-/-*^ BM pro-B cells and G1-arrested *Rag1*^*-/-*^ Wapl-degron *v-Abl* cells with or without Wapl depletion. Briefly, 10 million cells were collected and permeabilized with the DEPC treated buffer (10 mM Tris-HCl pH 7.4, 300 mM sucrose, 10 mM KCl, 5 mM MgCl2, 1 mM EGTA, 0.05% Tween-20, 0.1% NP40 substitute, 0.5 mM DTT, protease inhibitors and RNase inhibitor). The permeabilized cells were resuspended in 100 μl of DEPC treated storage buffer (10 mM Tris-HCl pH 8.0, 25% (v/v) glycerol, 5 mM MgCl2, 0.1 mM EDTA and 5 mM DTT) followed by nuclear run-on with 100 μl 2X run-on mix (5 mM Tris-HCl pH 8.0, 2.5 mM MgCl2, 0.5 mM DTT, 150mM KCl, 0.5 mM ATP, 0.5 mM CTP, 0.5 mM GTP, 0.5 mM Br-UTP, RNase inhibitor, 1% Sarkosyl) at 37 °C for 5 min. Total RNA was extracted by Trizol and followed by hydrolyzation with NaOH at a final concentration of 0.2 N on ice for 18 min. After quenching with ice-cold Tris-HCl pH 6.8 at a final concentration of 0.55 M and exchanging buffer via Bio-Rad P30 columns, the total RNA was incubated with Br-dU antibody-conjugated beads (Santa Cruz, #sc-32323-ac) for 1 h. The enriched Br-dU labeled RNAs were incubated with RppH (NEB, #M0356S) and with T4 PNK (NEB, #M0201S) for hydroxyl repair, followed by ligating the 5′ and 3′ RNA adaptors. RT-PCR was performed after adaptor ligation to obtain cDNAs. Half of the cDNAs was subjected to library preparation by two rounds of PCR with barcoded primers. The second round of PCR products were purified by AMPure beads (Beckman Coulter, #A63880). GRO-seq libraries were sequenced via paired-end 75 bp sequencing on Illumina NextSeq550. For comparison of all the GRO-seq results in this study, we used bowtie2 to align GRO-seq reads to mm9 genome and run MACS2 callpeak with parameters ‘—nomodel –keep-dup all -extsize 51 -nolambda -B -SPMR -g mm -broad’ to generate bigwig graph. To visualize the genome-wide RNA expression level, we run htseq-count to count read number on each gene and made scatter plot after down-sampling each sample to 10 million reads mappable to any genes. For comparison of GRO-seq results between WT and *Igh* inversion BM pro-B cells, GRO-seq profiles across the 4 V_H_ domains were potted in the normal orientation. Both the sense and antisense transcription are relative to the entire *Igh* V_H_ locus upstream V_H_81X with or without inversion and indicated respectively (Extended Data Fig. 3d). Spearman’s correlation coefficient (rho) and *p* values determined by two-sided Spearman’s correlation test are presented between indicated samples. *p* values are calculated and shown in the figures as the follows: non-significant (NS): *p*≥0.05, *: 0.01≤*p*<0.05, **: 0.001≤*p*<0.01, and ***: *p*<0.001.

### 3C-HTGTS and data analyses

3C-HTGTS on short-term cultured *Rag1*^*-/-*^ BM pro-B cells, G1-arrested Wapl-degron *v-Abl* cells without or with IAA&Dox treatment, was performed as previously described^10^. Briefly, 10 million cells were crosslinked with 2% formaldehyde (Sigma-Aldrich, #F8775) for 10 min at room temperature and quenched with glycine at a final concentration of 125 mM. Cells were lysed on ice for 10 min followed by centrifugation to get nuclei. Nuclei were resuspended in NEB Cutsmart buffer for NlaIII (NEB, #R0125) digestion at 37 °C overnight, followed by T4 DNA ligase (Promega, #M1801) mediated ligation under dilute conditions at 16 °C overnight. Ligated products were treated with Proteinase K (Roche, #03115852001) and RNase A (Invitrogen, #8003089) followed by DNA purification to get the 3C templates. 3C-HTGTS library preparation follows the standard LAM-HTGTS library preparation procedures as previously described^10^. 3C-HTGTS libraries were sequenced via Illumina NextSeq550 using paired-end 150-bp sequencing kit. For comparison, sequencing reads of all the 3C-HTGTS libraries were aligned to mm9 genome and processed as previously described^10^. Data were plotted for comparison after normalizing junction from each experimental 3C-HTGTS library by random selection to the total number of genome-wide or entire *Igh* locus (± 10 kb, chr12:114,444,081-117,258,165) junctions recovered from the smallest library in the set of libraries being compared. Chromosomal interaction patterns were very comparable before and after normalization. For short-term culture *Rag1*^*-/-*^ BM pro-B cells, 3C-HTGTS libraries using iEμ and V_H_8-7P (Fig. 1c, Extended Data Fig. 2) as baits were normalized to 885,651 and 742,577 total junctions of genome-wide, respectively. For comparison of iEμ-baited interactions between WT and *Igh* inversion BM pro-B cells in the inverted orientation, 3C-HTGTS results were inverted in the *Igh* inversion BM pro-B cells (Fig. 1c, bottom). For G1-arrested Wapl-degron *v-Abl* cells with or without Wapl depletion, 3C-HTGTS libraries using V_H_1-47 (Extended Data Fig. 7a) as baits were normalized to 231,542 total junctions of entire *Igh* locus (± 10 kb). For comparison of 3C-HTGTS libraries between short-term culture *Rag1*^*-/-*^ BM pro-B cells and G1-arrested Wapl-degron *v-Abl* cells with or without Wapl depletion, we removed the reads in the regions closely near to the baits (iEμ bait, chr12:114,664,385-114,668,233; V_H_8-7P bait, chr12:116,495,497-116,495,577), which did not contribute to the locus interaction. By this way, 3C-HTGTS libraries using iEμ (Fig. 3d; Extended Data Fig. 8d) and V_H_8-7P (Extended Data Fig. 7b) as baits were normalized to 53,795 and 195,259 total junctions of entire *Igh* locus (± 10 kb), respectively. The sequences of primers used for generating 3C-HTGTS libraries are listed in Supplementary Table 4.

### Data availability

HTGTS-V(D)J-Seq, 3C-HTGTS, ChIP-seq, and GRO-seq sequencing data reported in this study have been deposited in the GEO database under the accession number GSE151910. Specifically, HTGTS-V(D)J-Seq data related to Fig. 1b, 2, 3e-f and Extended Data Fig. 4, 8b-c, 8e-f, 9 are deposited in the GEO database under the accession number GSE151906. 3C-HTGTS data related to Fig. 1c, 3d, and Extended Data Fig. 2, 7, 8d are deposited in the GEO database under the accession number GSE151909. ChIP-seq data related to Extended Data Fig. 3a-c, 6a-b, 6e are deposited in the GEO database under the accession number GSE151908. GRO-seq data related to Fig. 3a-b and Extended Data Fig. 3d, 6c-d are deposited in the GEO database under the accession number GSE151907.

### Code availability

HTGTS-V(D)J-seq, 3C-HTGTS, ChIP-seq, and GRO-seq data were processed through the published pipelines as previously described^18^. Specifically, these pipelines are available at http://robinmeyers.github.io/transloc_pipeline/ (HTGTS pipeline), http://bowtie-bio.sourceforge.net/bowtie2/index.shtml (Bowtie2 v.2.2.8), https://sourceforge.net/projects/samtools/files/samtools/1.8/ (SAMtools v.1.8)

## Notes

### Competing Interest Statement

The authors have declared no competing interest.

