## Supplemental Data 1 for "Loop Extrusion Mediates Physiological Locus Contraction for V(D)J Recombination"

### Supplementary Data 1 I 3C-HTGTS, GRO-seq, ChIP-seq, CBE motif sites and PAIR elements for 15 representative peaks/clusters in $V_H$s domains in cultured *Rag1*$^{-/-}$ BM pro-B cells.

Related to Fig. 1c. Zoom-in profiles of 3C-HTGTS, GRO-seq, Rad21 ChIP-seq signals for ± 10 kb regions of 15 representative peaks/clusters in Fig.1c from *Rag1*$^{-/-}$ BM WT (blue) and *Igh* inversion (Red) pro-B cells along with relevant *bona fide* CBE motif sites were presented, PAIR elements (blue bars) and different $V_H$s that located in the same region were also shown above. For comparison, all the data were shown in normal *Igh* orientation.

From these results we could imply that the 2.4 Mb inversion didn't markedly disrupt $V_H$-locus contraction and major RC interactions with 15 representative major interaction points in the $V_H$ locus despite the wholesale change in the location and orientation of sequence across the locus (3C-HTGTS baiting from iEμ). In addition, germline $V_H$ transcription patterns (GRO-seq) and cohesion binding patterns (Rad21 ChIP-seq) across these 15 representative major interaction points were remained, largely as mirror images in the inverted locus.

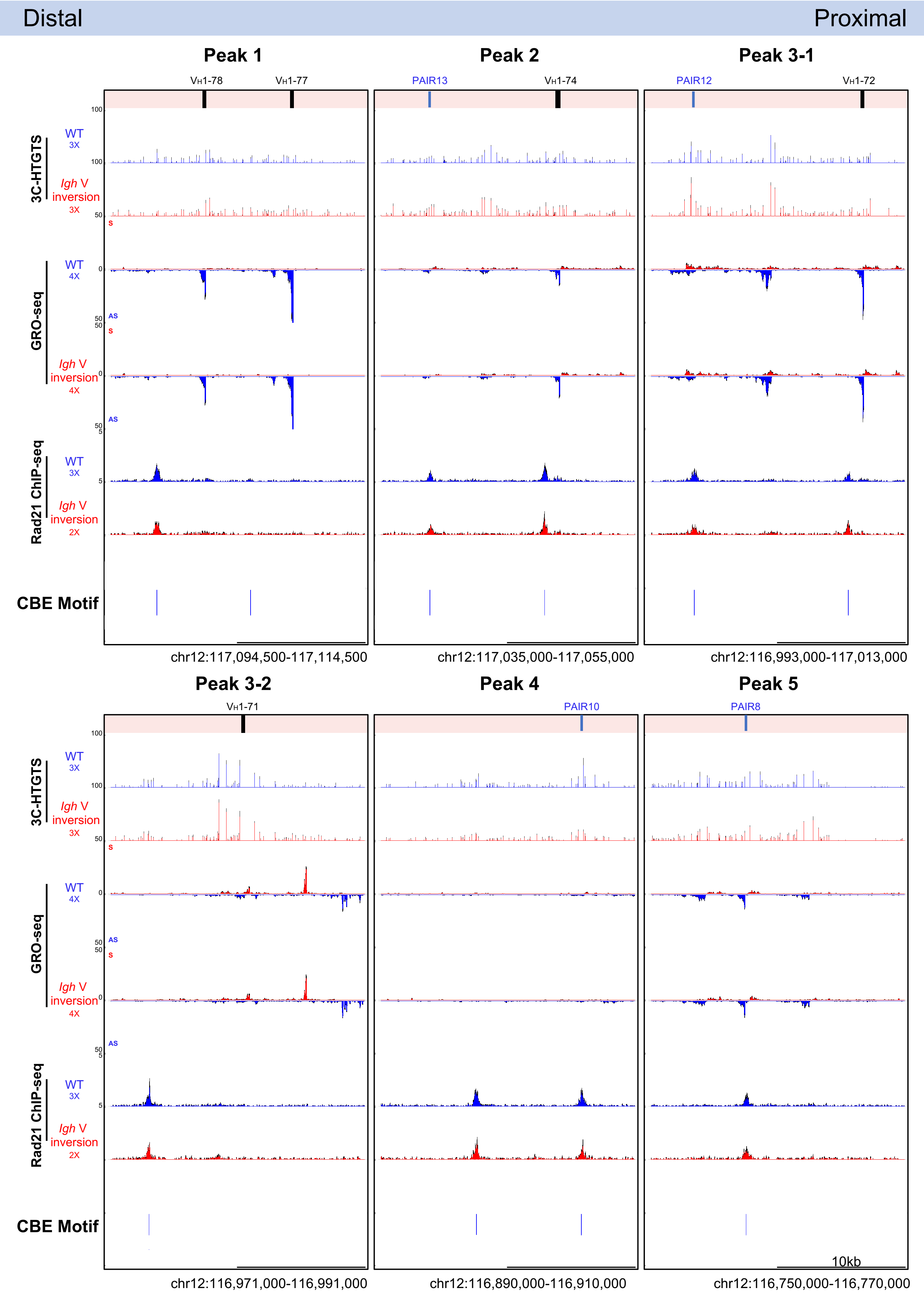

Distal
Proximal
Peak 1
Peak 2
Peak 3-1
Peak 3-2
Peak 4
Peak 5
VH1-78
VH1-77
PAIR13
VH1-74
PAIR12
VH1-72
VH1-71
PAIR10
PAIR8
3C-HTGTS
GRO-seq
Rad21 ChIP-seq
WT
Igh V inversion
3X
4X
2X
S
AS
CBE Motif
10kb
chr12:117,094,500-117,114,500
chr12:117,035,000-117,055,000
chr12:116,993,000-117,013,000
chr12:116,971,000-116,991,000
chr12:116,890,000-116,910,000
chr12:116,750,000-116,770,000

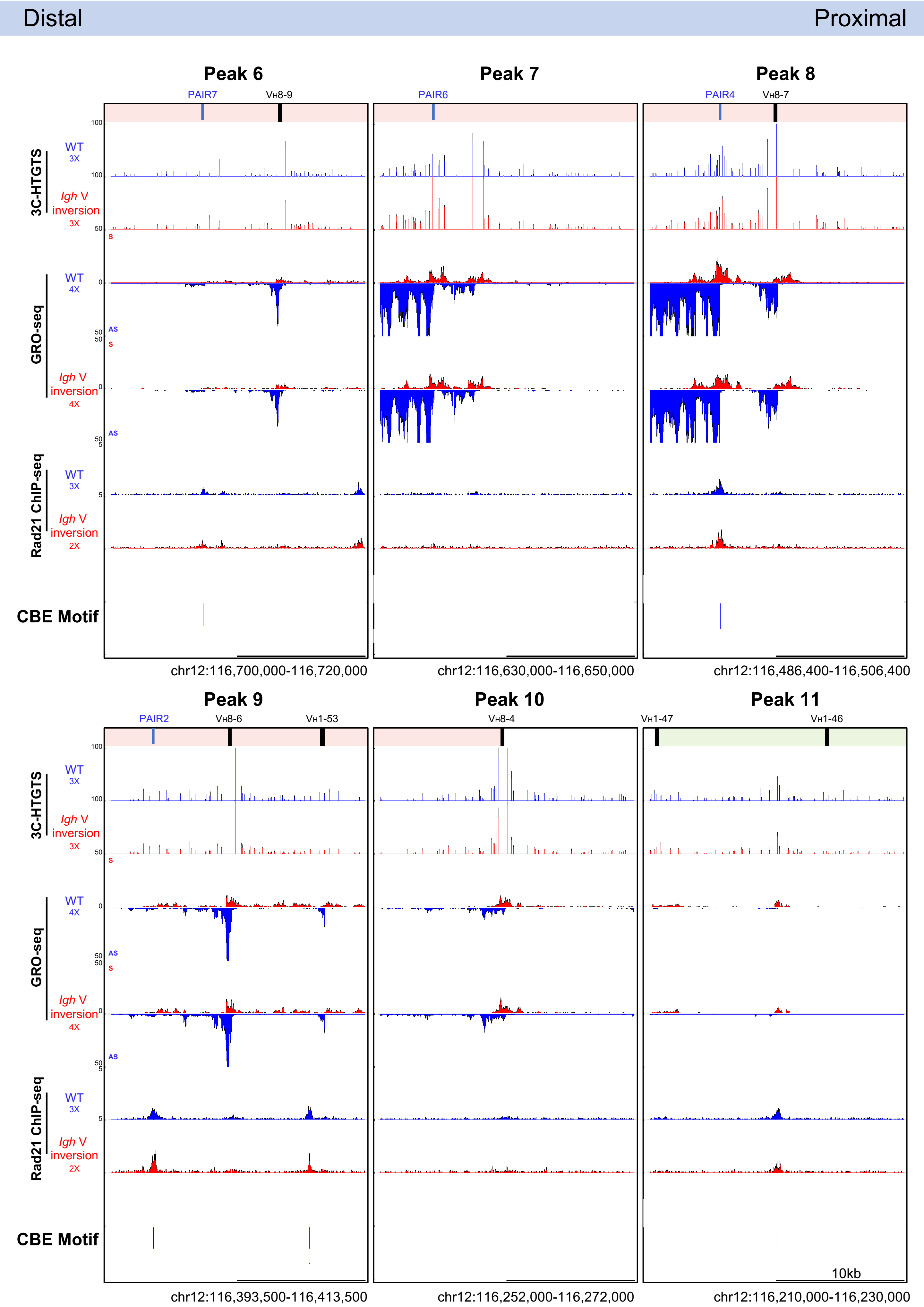

Distal
Proximal
Peak 6
Peak 7
Peak 8
PAIR7
VH8-9
PAIR6
PAIR4
VH8-7
3C-HTGTS
WT
3X
Igh V inversion
3X
GRO-seq
WT
4X
Igh V inversion
4X
Rad21 ChIP-seq
WT
3X
Igh V inversion
2X
CBE Motif
chr12:116,700,000-116,720,000
chr12:116,630,000-116,650,000
chr12:116,486,400-116,506,400
Peak 9
Peak 10
Peak 11
PAIR2
VH8-6
VH1-53
VH8-4
VH1-47
VH1-46
10kb
chr12:116,393,500-116,413,500
chr12:116,252,000-116,272,000
chr12:116,210,000-116,230,000

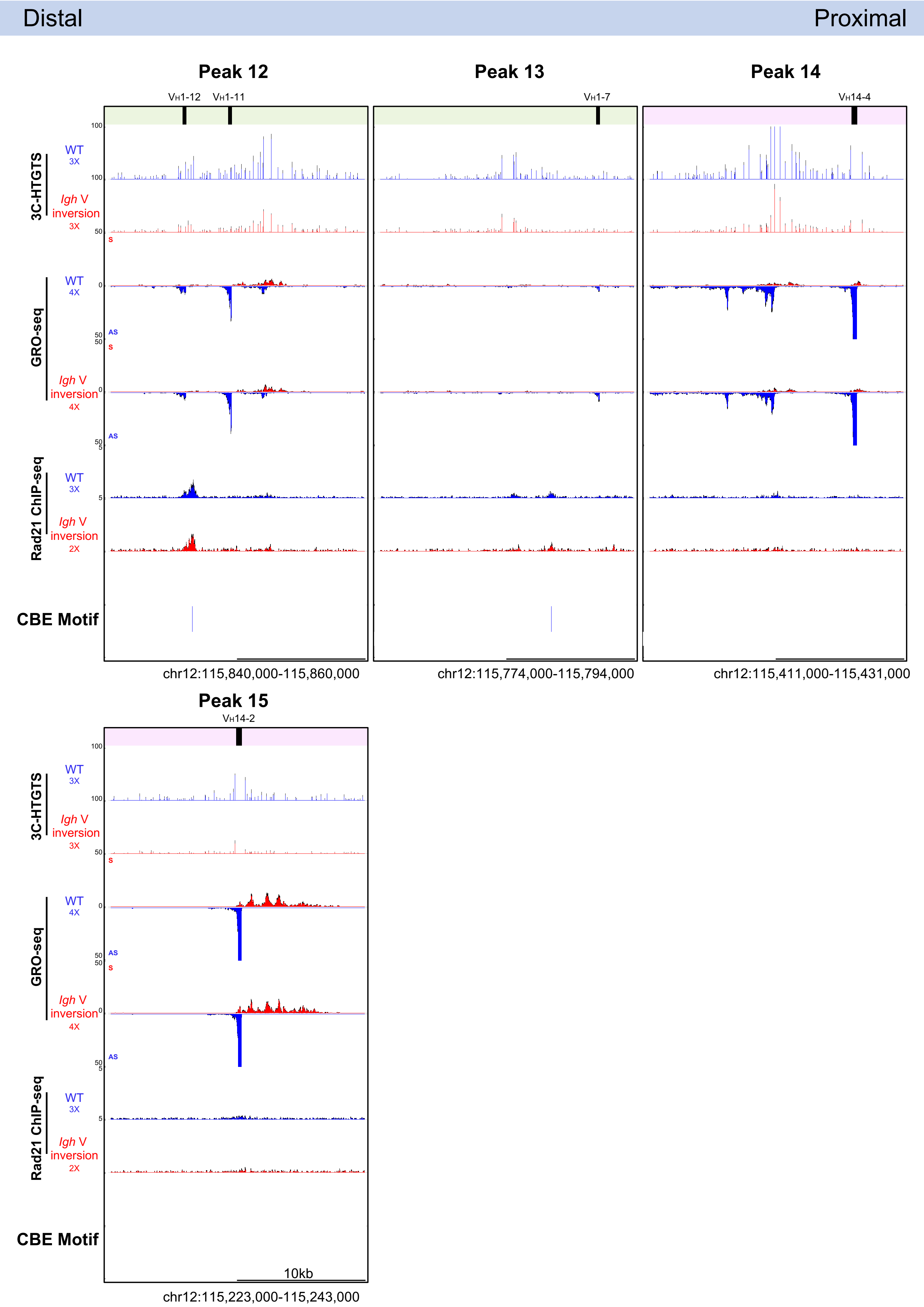

Distal
Proximal
Peak 12
VH1-12
VH1-11
Peak 13
VH1-7
Peak 14
VH14-4
3C-HTGTS
WT
3X
Igh V inversion
3X
GRO-seq
WT
4X
Igh V inversion
4X
Rad21 ChIP-seq
WT
3X
Igh V inversion
2X
CBE Motif
100
50
0
5
S
AS
chr12:115,840,000-115,860,000
chr12:115,774,000-115,794,000
chr12:115,411,000-115,431,000
Peak 15
VH14-2
10kb
chr12:115,223,000-115,243,000
