## Supplemental Data 2 for "Loop Extrusion Mediates Physiological Locus Contraction for V(D)J Recombination"

### Supplementary Data 2 l $V_H$ usage, GRO-seq, 3C-HTGTS, ChIP-seq, CBE motifs sites and PAIR elements for all $V_H$s in G1-arrested Wapl-degron *v-Abl* pro-B cells.

Average $V_H$ usage ± s.d. and zoom-in profile of GRO-seq, 3C-HTGTS, Rad-21/CTCF ChIP-seq signals for ± 10kb region of all $V_H$s in G1-arrested Wapl degron *v-Abl* line without (Untreated, blue) or with (IAA&Dox, red) Wapl depletion along with relevant *bona fide* CBE motif sites are presented, PAIR elements (blue bars) that located near indicated $V_H$ were also shown above.

Utilization data of $V_H$ segments was normalized to 100,000 total recovered junctions. We used unpaired two-tailed t-test to calculated the *p* value. NA, not applicable, which means average read<1; p ≥0.05 means NS: no significant; *0.01≤p<0.05, **0.001≤p<0.01, ***p<0.001.

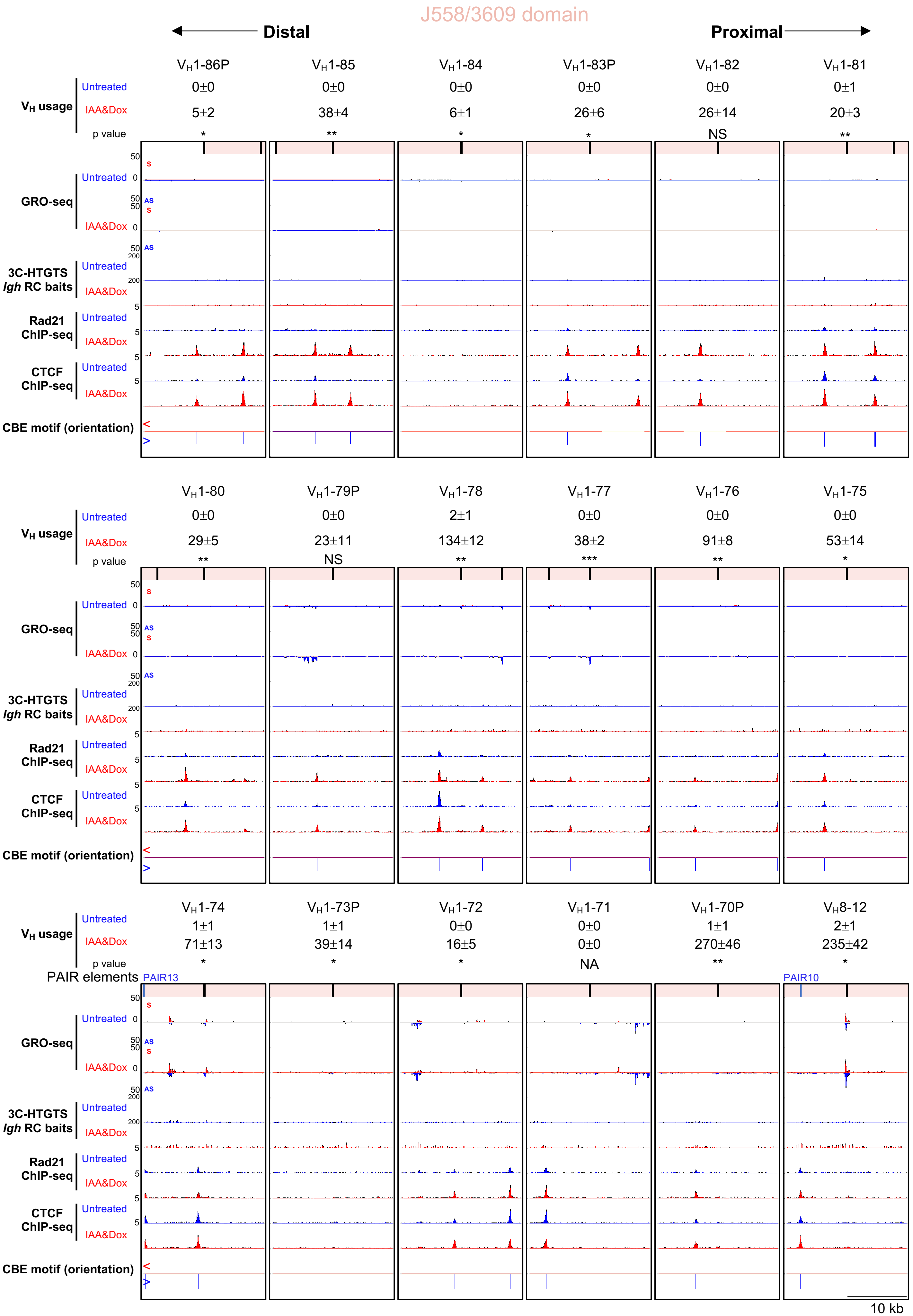

J558/3609 domain
Distal
Proximal
$V_H$ usage
Untreated
IAA&Dox
p value
$V_H$1-86P 0±0 5±2 *
$V_H$1-85 0±0 38±4 **
$V_H$1-84 0±0 6±1 *
$V_H$1-83P 0±0 26±6 *
$V_H$1-82 0±0 26±14 NS
$V_H$1-81 0±1 20±3 **
$V_H$1-80 0±0 29±5 **
$V_H$1-79P 0±0 23±11 NS
$V_H$1-78 2±1 134±12 **
$V_H$1-77 0±0 38±2 ***
$V_H$1-76 0±0 91±8 **
$V_H$1-75 0±0 53±14 *
$V_H$1-74 1±1 71±13 *
$V_H$1-73P 1±1 39±14 *
$V_H$1-72 0±0 16±5 *
$V_H$1-71 0±0 0±0 NA
$V_H$1-70P 1±1 270±46 **
$V_H$8-12 2±1 235±42 *
GRO-seq
3C-HTGTS
*Igh* RC baits
Rad21 ChIP-seq
CTCF ChIP-seq
CBE motif (orientation)
PAIR elements
PAIR13
PAIR10
S
AS
10 kb

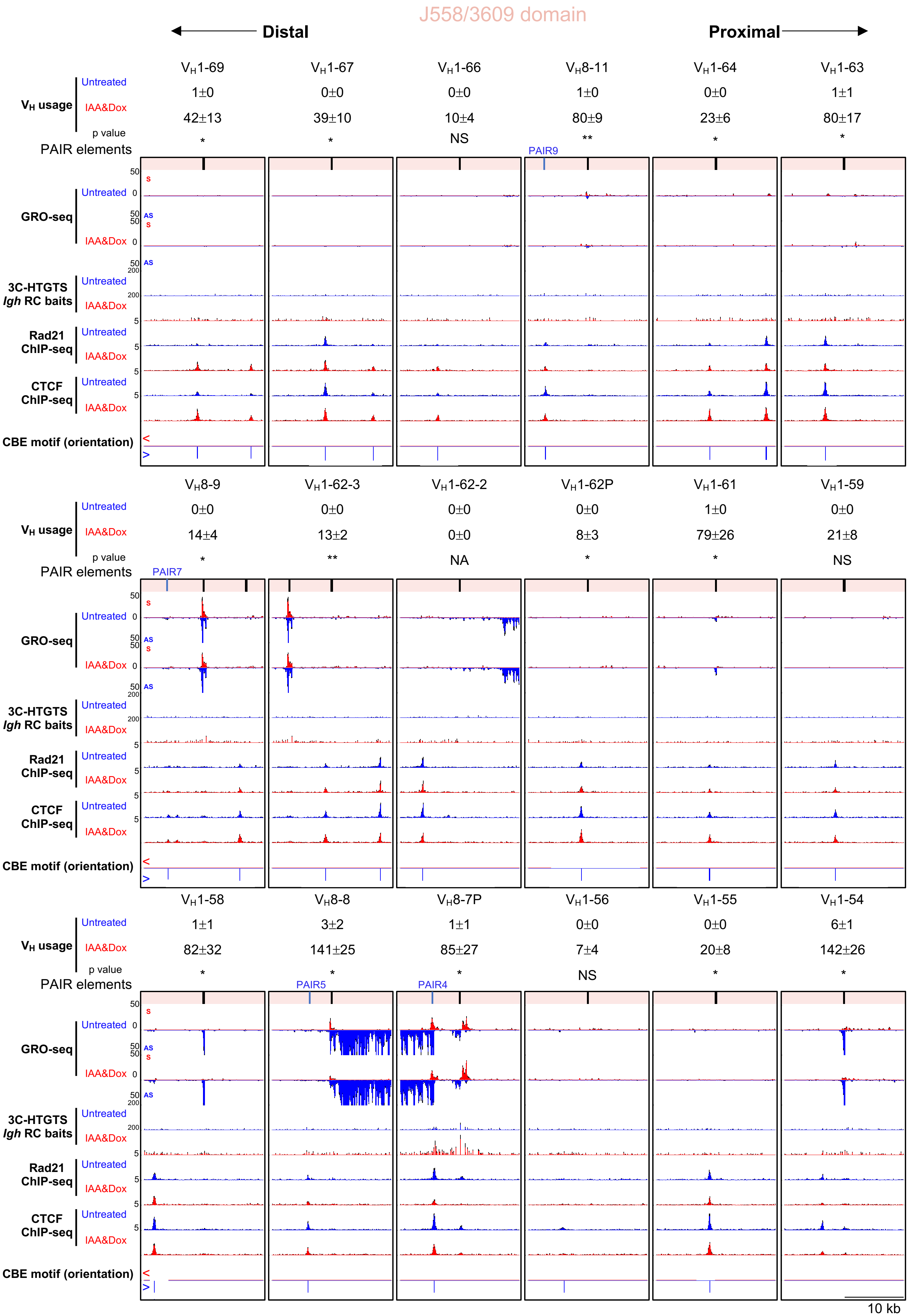

J558/3609 domain
Distal
Proximal
VH usage
Untreated
IAA&Dox
p value
PAIR elements
GRO-seq
3C-HTGTS
Igh RC baits
Rad21
ChIP-seq
CTCF
ChIP-seq
CBE motif (orientation)
VH1-69 | 1±0 | 42±13 | *
VH1-67 | 0±0 | 39±10 | *
VH1-66 | 0±0 | 10±4 | NS
VH8-11 | 1±0 | 80±9 | **
VH1-64 | 0±0 | 23±6 | *
VH1-63 | 1±1 | 80±17 | *
PAIR9
VH8-9 | 0±0 | 14±4 | *
VH1-62-3 | 0±0 | 13±2 | **
VH1-62-2 | 0±0 | 0±0 | NA
VH1-62P | 0±0 | 8±3 | *
VH1-61 | 1±0 | 79±26 | *
VH1-59 | 0±0 | 21±8 | NS
PAIR7
VH1-58 | 1±1 | 82±32 | *
VH8-8 | 3±2 | 141±25 | *
VH8-7P | 1±1 | 85±27 | *
VH1-56 | 0±0 | 7±4 | NS
VH1-55 | 0±0 | 20±8 | *
VH1-54 | 6±1 | 142±26 | *
PAIR5
PAIR4
S
AS
50
0
200
5
10 kb

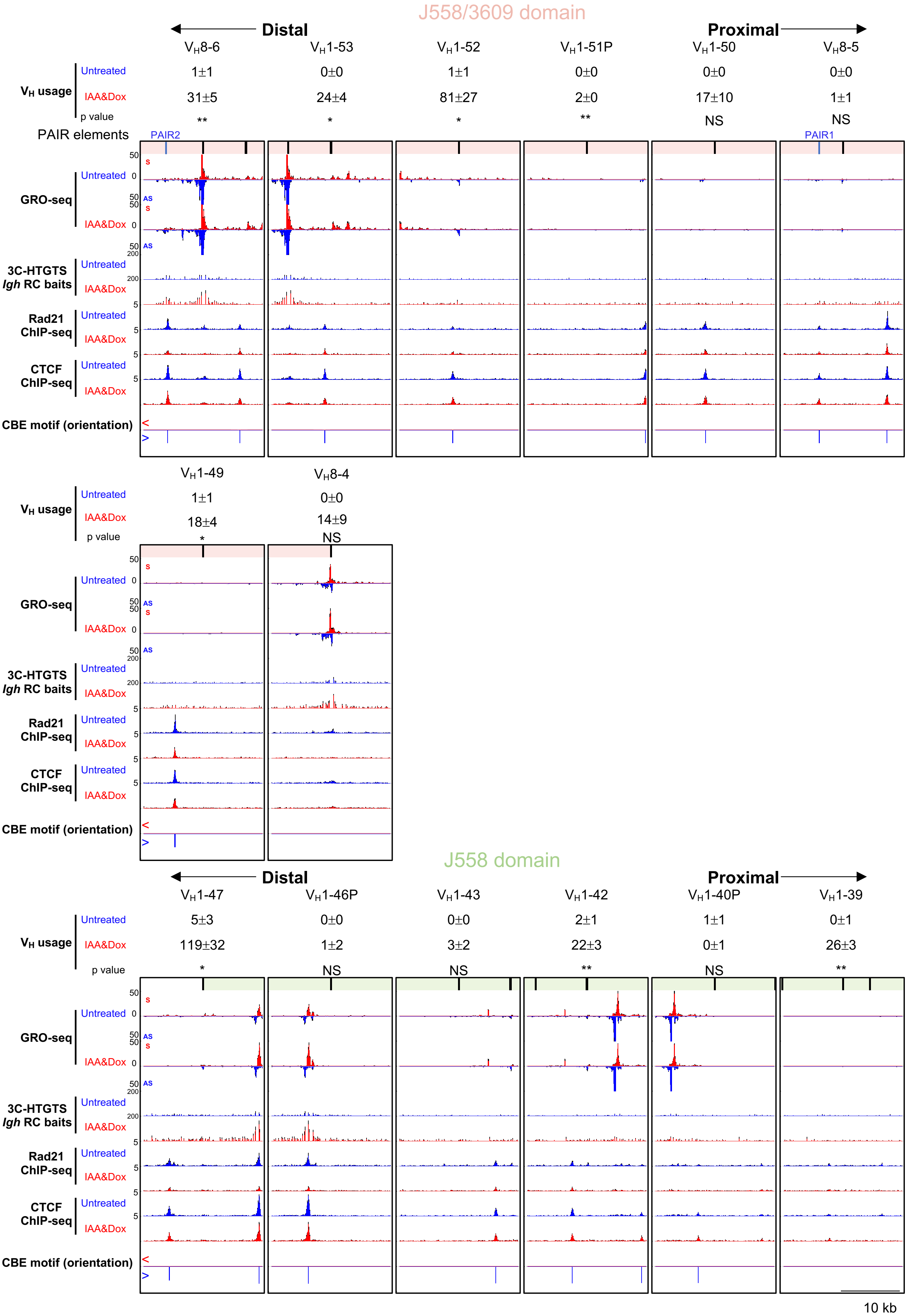
J558/3609 domain
Distal
Proximal
$V_H$8-6
$V_H$1-53
$V_H$1-52
$V_H$1-51P
$V_H$1-50
$V_H$8-5
$V_H$ usage
Untreated
1±1
0±0
1±1
0±0
0±0
0±0
IAA&Dox
31±5
24±4
81±27
2±0
17±10
1±1
p value
**
*
*
**
NS
NS
PAIR elements
PAIR2
PAIR1
GRO-seq
3C-HTGTS
*Igh* RC baits
Rad21
ChIP-seq
CTCF
ChIP-seq
CBE motif (orientation)
$V_H$1-49
$V_H$8-4
1±1
0±0
18±4
14±9
*
NS
J558 domain
$V_H$1-47
$V_H$1-46P
$V_H$1-43
$V_H$1-42
$V_H$1-40P
$V_H$1-39
5±3
0±0
0±0
2±1
1±1
0±1
119±32
1±2
3±2
22±3
0±1
26±3
*
NS
NS
**
NS
**
10 kb

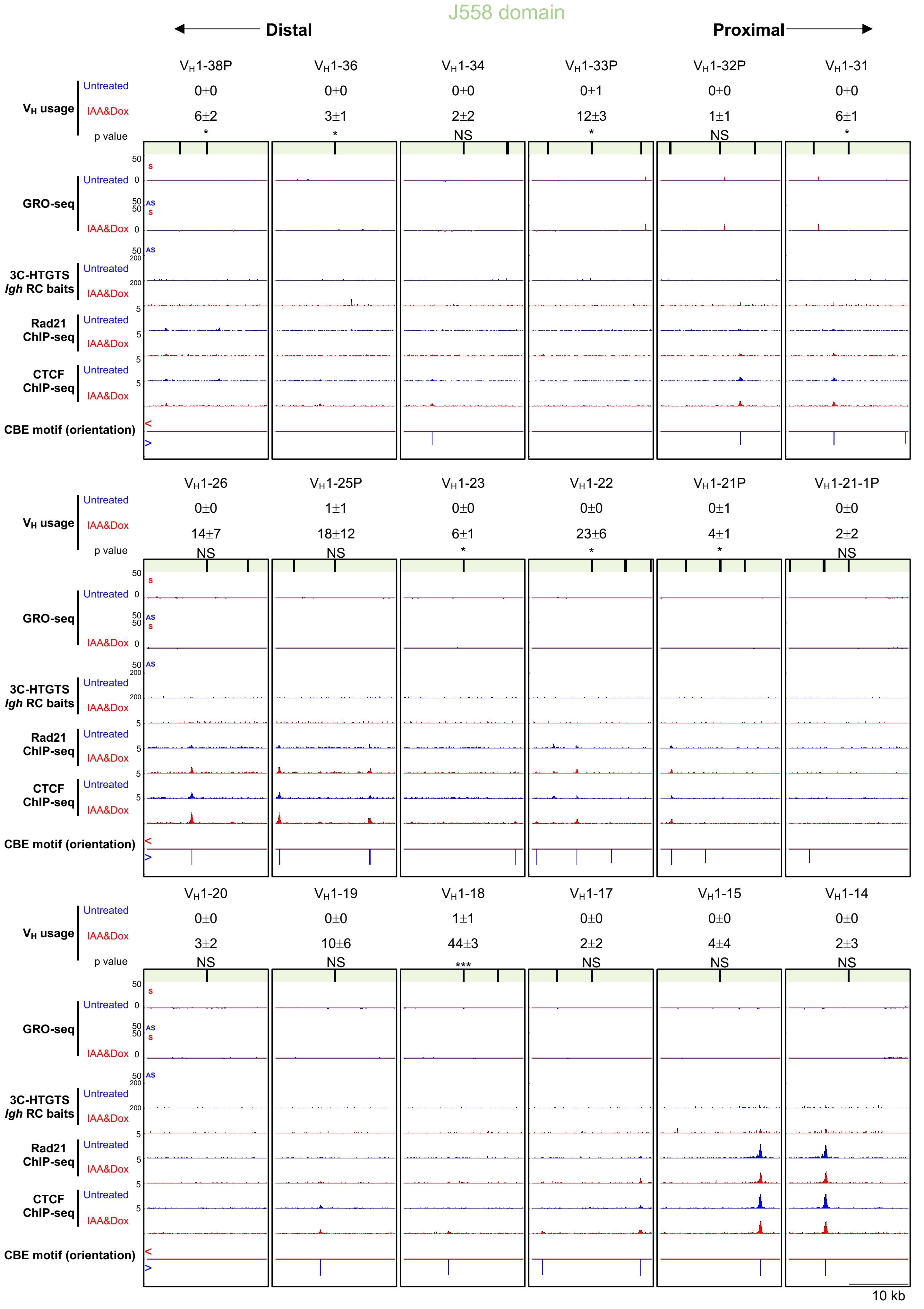
J558 domain
Distal
Proximal
VH1-38P
VH1-36
VH1-34
VH1-33P
VH1-32P
VH1-31
VH usage
Untreated
IAA&Dox
p value
0±0
0±0
0±0
0±1
0±0
0±0
6±2
3±1
2±2
12±3
1±1
6±1
*
*
NS
*
NS
*
GRO-seq
3C-HTGTS
Igh RC baits
Rad21 ChIP-seq
CTCF ChIP-seq
CBE motif (orientation)
VH1-26
VH1-25P
VH1-23
VH1-22
VH1-21P
VH1-21-1P
0±0
1±1
0±0
0±0
0±1
0±0
14±7
18±12
6±1
23±6
4±1
2±2
NS
NS
*
*
*
NS
VH1-20
VH1-19
VH1-18
VH1-17
VH1-15
VH1-14
0±0
0±0
1±1
0±0
0±0
0±0
3±2
10±6
44±3
2±2
4±4
2±3
NS
NS
***
NS
NS
NS
10 kb

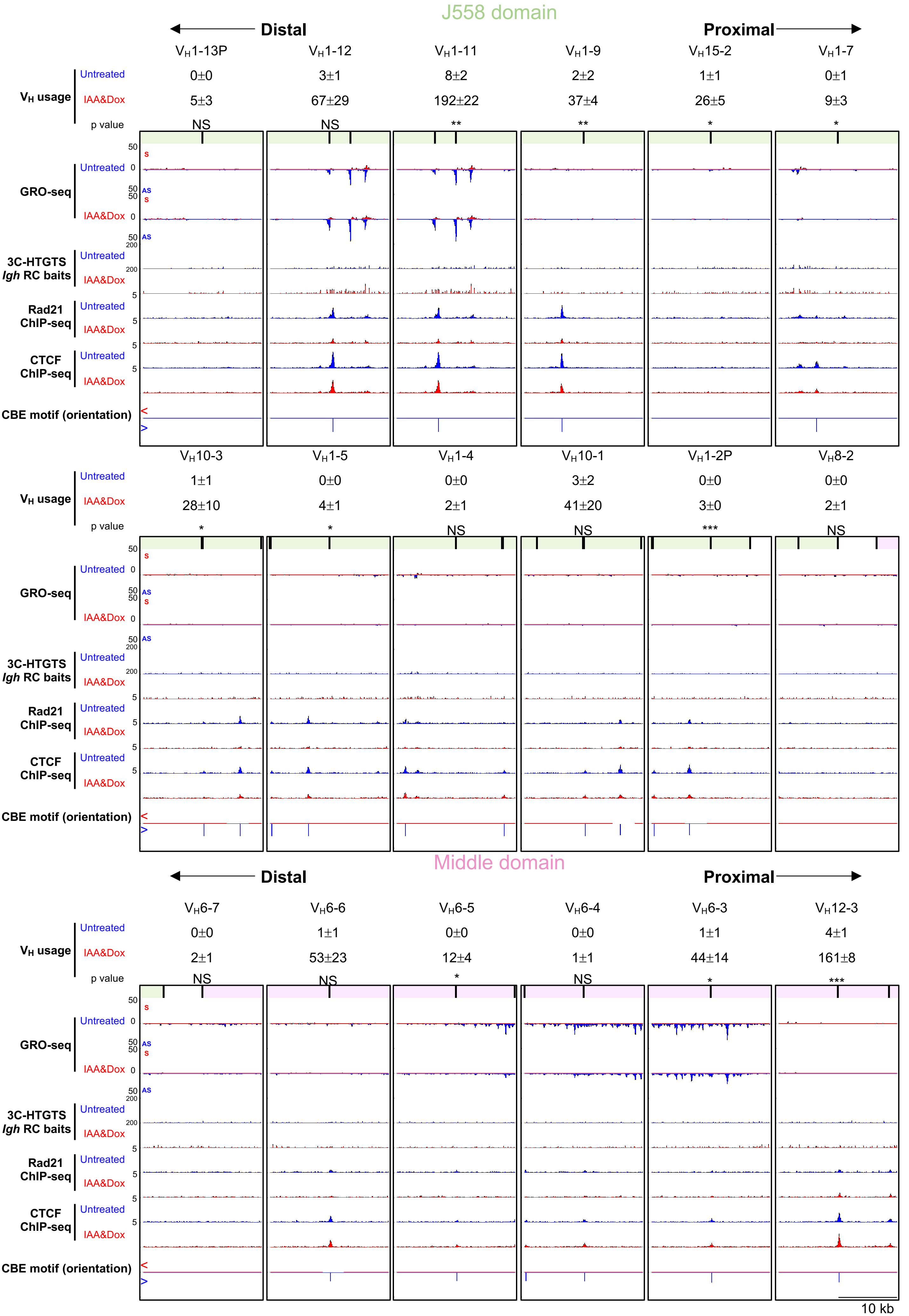

J558 domain
Distal
Proximal
$V_H$1-13P
$V_H$1-12
$V_H$1-11
$V_H$1-9
$V_H$15-2
$V_H$1-7
$V_H$ usage
Untreated
0±0
3±1
8±2
2±2
1±1
0±1
IAA&Dox
5±3
67±29
192±22
37±4
26±5
9±3
p value
NS
NS
**
**
*
*
GRO-seq
S
AS
3C-HTGTS
Igh RC baits
Rad21 ChIP-seq
CTCF ChIP-seq
CBE motif (orientation)
$V_H$10-3
$V_H$1-5
$V_H$1-4
$V_H$10-1
$V_H$1-2P
$V_H$8-2
Untreated
1±1
0±0
0±0
3±2
0±0
0±0
IAA&Dox
28±10
4±1
2±1
41±20
3±0
2±1
p value
*
*
NS
NS
***
NS
Middle domain
Distal
Proximal
$V_H$6-7
$V_H$6-6
$V_H$6-5
$V_H$6-4
$V_H$6-3
$V_H$12-3
Untreated
0±0
1±1
0±0
0±0
1±1
4±1
IAA&Dox
2±1
53±23
12±4
1±1
44±14
161±8
p value
NS
NS
*
NS
*
***
10 kb

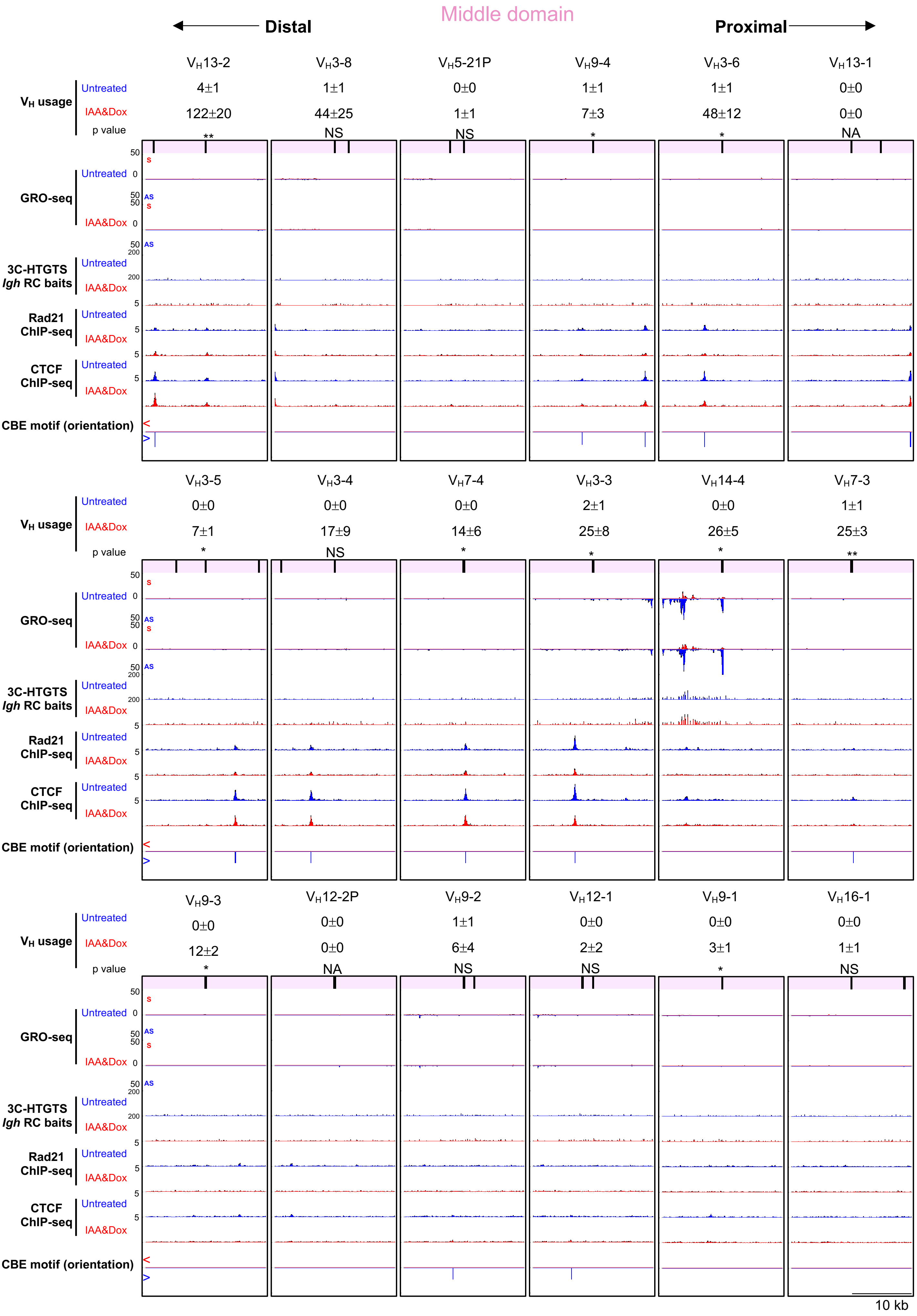
Middle domain
Distal
Proximal
V_H13-2 V_H3-8 V_H5-21P V_H9-4 V_H3-6 V_H13-1
V_H usage
Untreated 4±1 1±1 0±0 1±1 1±1 0±0
IAA&Dox 122±20 44±25 1±1 7±3 48±12 0±0
p value ** NS NS * * NA
V_H3-5 V_H3-4 V_H7-4 V_H3-3 V_H14-4 V_H7-3
Untreated 0±0 0±0 0±0 2±1 0±0 1±1
IAA&Dox 7±1 17±9 14±6 25±8 26±5 25±3
p value * NS * * * **
V_H9-3 V_H12-2P V_H9-2 V_H12-1 V_H9-1 V_H16-1
Untreated 0±0 0±0 1±1 0±0 0±0 0±0
IAA&Dox 12±2 0±0 6±4 2±2 3±1 1±1
p value * NA NS NS * NS
GRO-seq
Untreated
IAA&Dox
S
AS
50
0
3C-HTGTS
Igh RC baits
200
Rad21 ChIP-seq
CTCF ChIP-seq
5
CBE motif (orientation)
<
>
10 kb

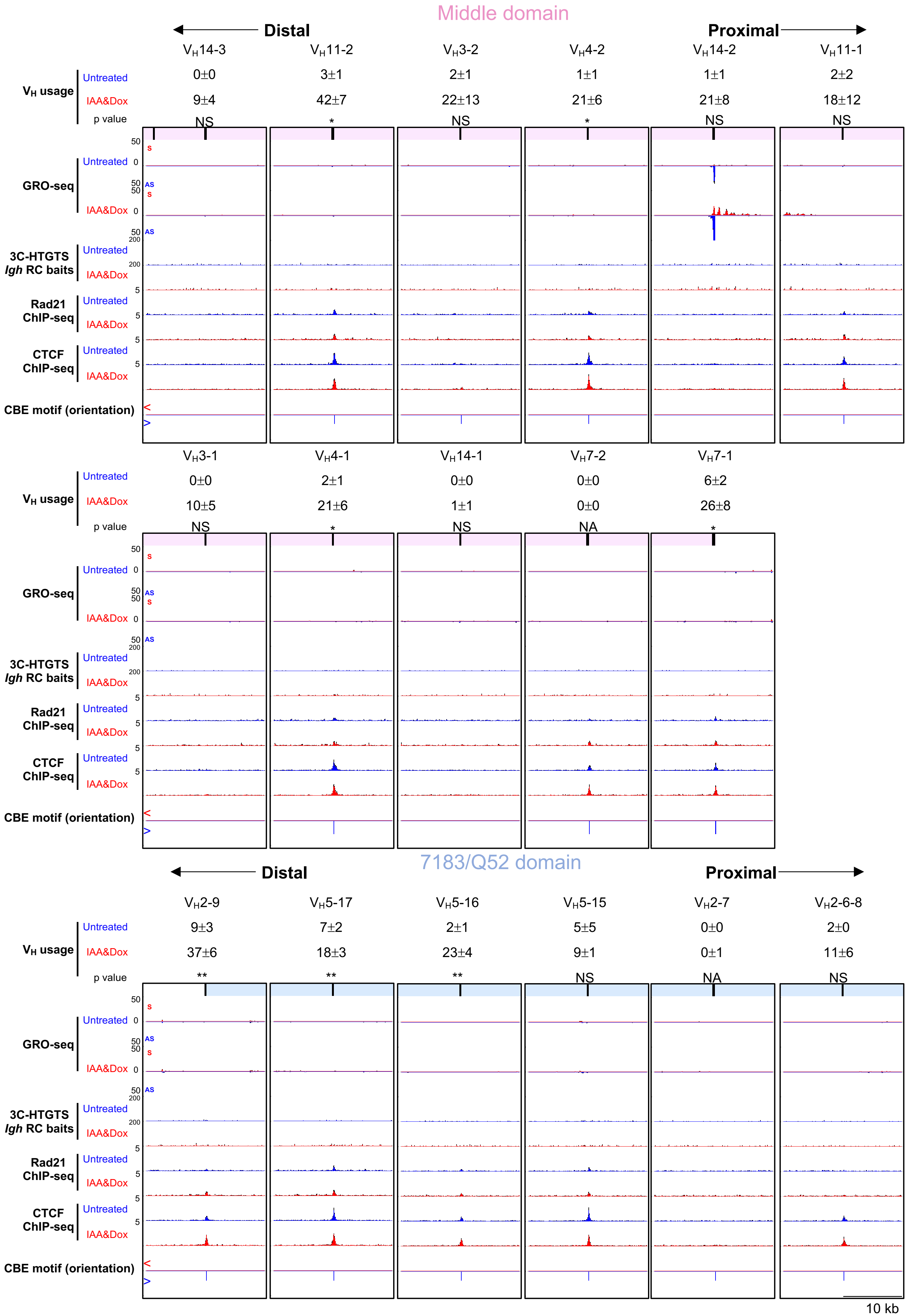
Middle domain
Distal
Proximal
VH14-3 VH11-2 VH3-2 VH4-2 VH14-2 VH11-1
VH usage
Untreated 0±0 3±1 2±1 1±1 1±1 2±2
IAA&Dox 9±4 42±7 22±13 21±6 21±8 18±12
p value NS * NS * NS NS
GRO-seq
3C-HTGTS
Igh RC baits
Rad21 ChIP-seq
CTCF ChIP-seq
CBE motif (orientation)
VH3-1 VH4-1 VH14-1 VH7-2 VH7-1
Untreated 0±0 2±1 0±0 0±0 6±2
IAA&Dox 10±5 21±6 1±1 0±0 26±8
p value NS * NS NA *
7183/Q52 domain
VH2-9 VH5-17 VH5-16 VH5-15 VH2-7 VH2-6-8
Untreated 9±3 7±2 2±1 5±5 0±0 2±0
IAA&Dox 37±6 18±3 23±4 9±1 0±1 11±6
p value ** ** ** NS NA NS
10 kb

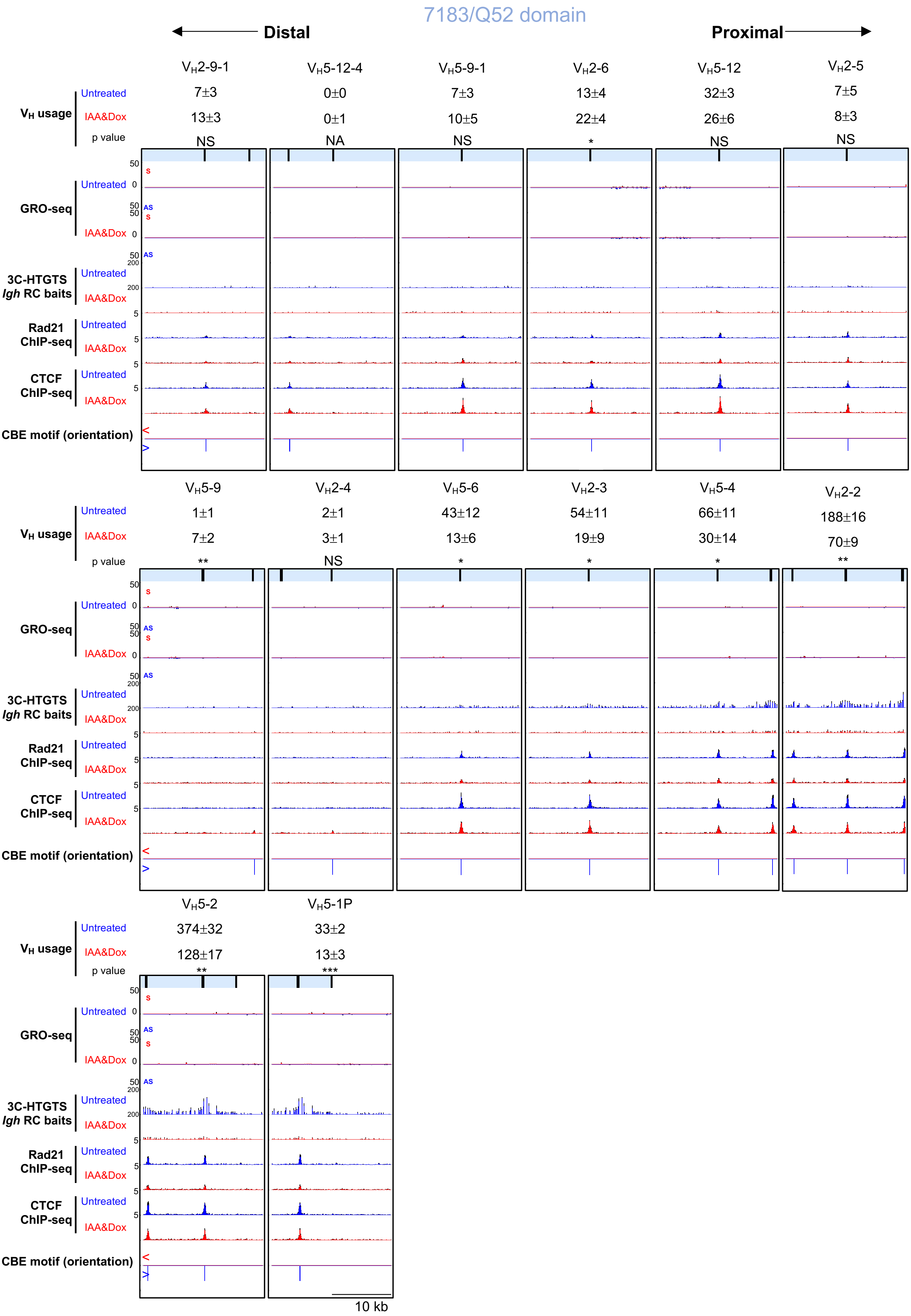
7183/Q52 domain
Distal
Proximal
VH2-9-1
VH5-12-4
VH5-9-1
VH2-6
VH5-12
VH2-5
VH usage
Untreated
7±3
0±0
7±3
13±4
32±3
7±5
IAA&Dox
13±3
0±1
10±5
22±4
26±6
8±3
p value
NS
NA
NS
*
NS
NS
GRO-seq
S
AS
3C-HTGTS
Igh RC baits
Rad21
ChIP-seq
CTCF
ChIP-seq
CBE motif (orientation)
VH5-9
VH2-4
VH5-6
VH2-3
VH5-4
VH2-2
1±1
2±1
43±12
54±11
66±11
188±16
7±2
3±1
13±6
19±9
30±14
70±9
**
NS
*
*
*
**
VH5-2
VH5-1P
374±32
33±2
128±17
13±3
**
***
10 kb
